# A computable probability landscape for T cell receptor repertoire sampling and comparison

**DOI:** 10.64898/2026.10.15.759459

**Authors:** Thomas Konstantinovsky, Gur Yaari

## Abstract

Finite sampling complicates interpretation and comparison of T cell receptor (TCR) repertoires. We introduce BEAM, a probability model for TCR*β* complementarity-determining region 3 amino-acid sequences that combines local sequence dependence with exact normalisation. Its tractable local structure permits deterministic evaluation of sequence-pattern probabilities and sampling expectations without enumerating or Monte Carlo sampling the sequence domain. A reference built from 38,538 human repertoires links sampling depth to expected sequence recovery, recurrence, overlap and publicness, the number of cohort repertoires containing a sequence. Models built from individual repertoires define distances attributable to sequence lengths and local patterns. BEAM improves held-out sequence prediction over controls matched for length and individual-position frequencies, ranks sequence publicness in an external cohort and matches CD4 and CD8 repertoires across independent studies. BEAM therefore provides a tractable observational probability landscape for depth-aware sampling calculations and interpretable repertoire comparison.

---

T cell receptor (TCR) sequencing reveals immune history and genetic background [1, 2], but finite and uneven sampling complicates interpreting distinct-sequence counts, sharing and repertoire differences [3, 4, 5]. Publicness, the number of repertoires in a cohort containing a sequence, depends on both biology and sampling depth. Sequences found in only one cohort member are private, making both labels specific to the cohort examined. The complementarity-determining region 3 of the TCR*β* chain (CDR3*β*) contributes to antigen specificity; its observed distribution reflects recombination, selection and clonal expansion [6]. Repeated amino-acid sequences can arise from expansion or independent rearrangements [7, 8]. Interpreting recurrence, sharing and repertoire differences therefore requires probabilistic sampling baselines.

Existing models provide complementary baselines. Models of variable, diversity and joining (V(D)J) gene recombination infer generation probabilities, *P*_gen_ [9, 10]; OLGA computes these efficiently for amino-acid sequences [11]. SoNIA and soNNia reweight generation probabilities to describe selection in observed repertoires [12, 13, 14]. These models predict sharing, although whole-space sampling quantities are typically evaluated using generated sequences [4]. LZGraphs learn recurring subsequences directly [15]. For repertoire comparison, gene-usage measures and receptor-similarity methods capture different aspects of empirical samples [5, 16, 17]; fitted selection models have also been compared as probability distributions [13]. A tractable observational model could make global sampling expectations directly computable while supporting repertoire comparison through sequence patterns represented in the fitted distributions, including those absent from finite samples.

Here we introduce BEAM (Bidirectional Expert-Anchored Model), which learns local aminoacid relationships from sequence counts using contexts anchored at both CDR3*β* ends (Fig. 1). End-relative indexing shares parameters across lengths [13], and hierarchical smoothing supplies probabilities for unseen contexts. The local structure permits exact normalisation and marginal probabilities by dynamic programming. Expected distinct-sequence counts, recurrence, overlap and publicness can then be evaluated numerically without enumerating or sampling the model’s sequence domain (Methods 3.2, 3.5).

**Figure 1:**
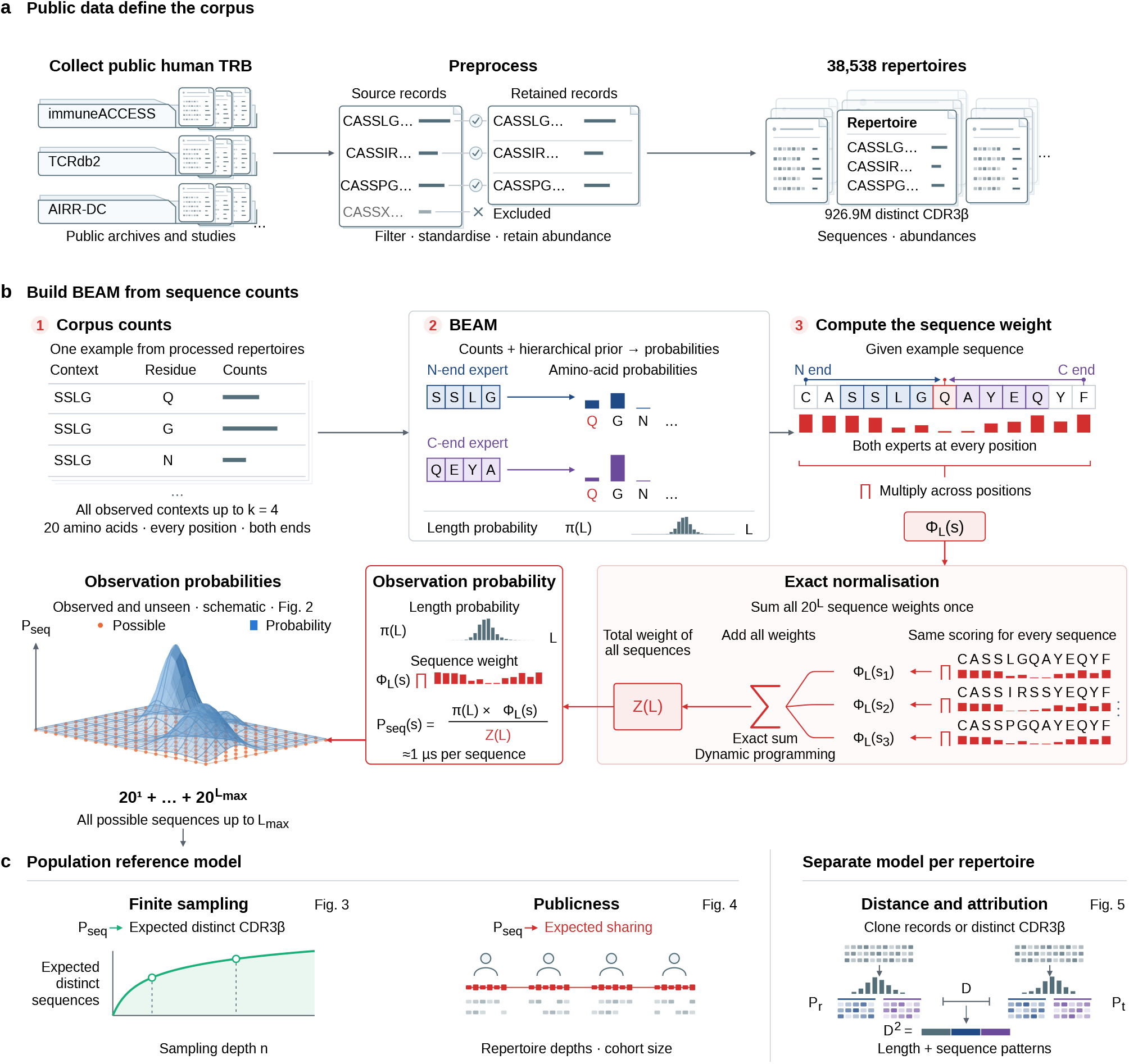
BEAM: from public repertoires to observation probabilities and repertoire models. (**a**) Public human TRB repertoires are filtered to CDR3*β* strings of 3–27 standard residues; non-standard strings are excluded ( *×* ). Platform-reported template counts are retained where available; otherwise each distinct amino-acid sequence receives unit weight within its repertoire. The corpus contains 38,538 repertoires and 926,909,993 pooled distinct sequences. (**b**) Model construction. (1) Residue counts by contexts of up to 4 residues and position from either end; count bars are schematic. (2) N-end and C-end experts convert counts into probabilities using a hierarchical prior; *π*(L) gives length probability. Blue and purple bars show population reference model expert-row probabilities for Q, G and N among 20 residues, after SSLG from the N end and QEYA from the C end, respectively. (3) Red bars show combined residue weights for one sequence; their product gives Φ_*L*_(s). Dynamic programming sums weights over all 20^*L*^ strings without listing them, yielding exact normalisation Z(L). Three length-13 sequences illustrate this sum. The sequence observation probability is *P*_seq_(s) = *π*(L) Φ_*L*_(s) / Z(L); scoring averages 1 *µ*s per sequence with the model loaded. The schematic landscape represents observation probabilities for observed and unseen strings. The population reference model’s domain is all standard-amino-acid strings of lengths 1–27 (1.41 *×* 10^35^ strings), rather than a count of biologically realisable receptors. (**c**) Two model-use branches. **Population reference model:** independent sampling at specified repertoire depths and cohort size predicts distinct sequence counts (Fig. 3) and publicness (Fig. 4). **Separate model per repertoire:** each repertoire supplies counts for its own BEAM model, with the input units specified in Fig. 5. Their local-block Hellinger distance D compares repertoires; D^2^ is attributed to length and sequence-pattern contributions (Fig. 5). The small contribution bar is schematic; segment widths are illustrative.

We fitted a population reference to 926,909,993 distinct CDR3*β* sequences from38,538human repertoires, using template counts where available and unit weight per distinct sequence within each remaining repertoire (Methods 3.1). Observation probability, *P*_seq_, denotes the probability of a sequence on one draw from the fitted distribution; it combines the biological and measurement processes represented in its inputs. The same procedure fits separate repertoire models without gene annotations. Population reference models support sampling and publicness calculations; perrepertoire models support distance and attribution to sequence features.

We test held-out sequence prediction beyond length and positional composition (Fig. 2e), evaluate depth-dependent sequence accumulation (Fig. 3) and examine publicness across cohorts (Fig. 4). We then assess repertoire distances across sorted populations and independent human studies and trace their contributions to interpretable local sequence features (Fig. 5). BEAM thus connects fitted observation probabilities to quantitative sampling baselines and interpretable repertoire comparisons.

**Figure 2:**
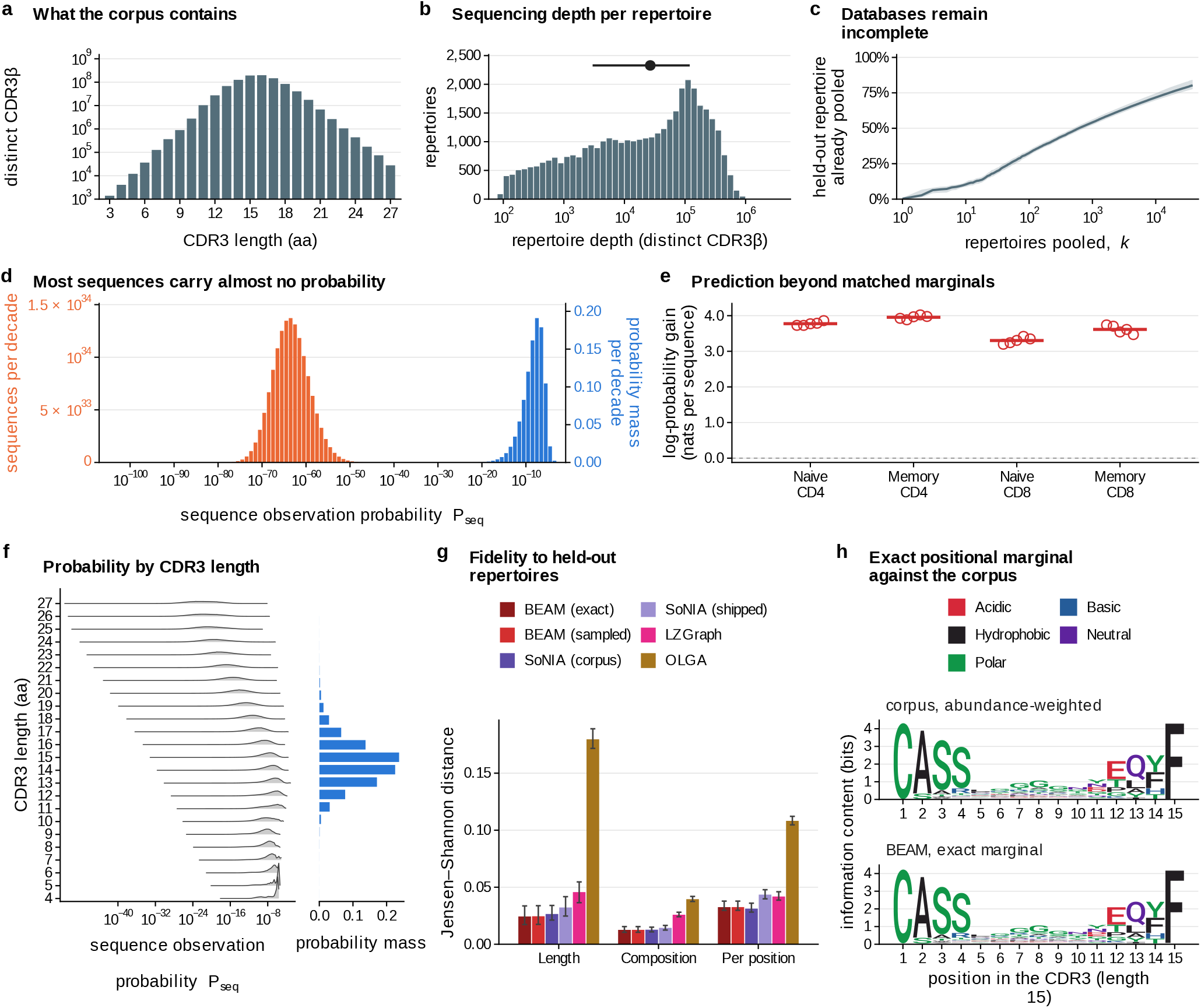
The corpus, probability landscape and held-out sequence prediction. (**a**) Distinct retained CDR3*β* by length: 926,909,993 sequences from 38,538 repertoires; logarithmic scale. (**b**) Distinct CDR3*β* per repertoire (0.1-decade bins; logarithmic x axis); point and segment, median and interquartile range. (**c**) Fraction of a held-out repertoire’s sequences found in k pooled repertoires (logarithmic k axis). 300 repertoires were held out; 38,238 accumulated in 10 random orders. Line and band, median and interquartile range over held-out repertoires and orders. (**d**) Discretised spectrum per decade of sequence observation probability *P*_seq_. Orange, left axis: number of possible standard-amino-acid strings of 1–27 residues. Blue, right axis: probability mass. Paired bars split each decade; linear axes are rescaled to align maxima. (**e**) Mean natural-log probability gain over Q, the independent-position projection matching the same BEAM fit’s length distribution and every within-length positional marginal. Each of 5 Rubelt twin families [18] is held out in turn; separate models for 4 populations use the other 8 donors. Test strings are absent from all training populations. Open points average the two held-out donor means; horizontal bars average families. Input records have unit weight per nucleotide-defined clone key. Positive gain means lower cross-entropy; dashed line, zero. Training folds overlap. (**f** ) Probability-weighted *P*_seq_ distributions by length, from 4 residues: unit-area densities (left); probability mass per length (right). (**g**) Jensen–Shannon distance from 768 held-out Emerson donors [1] for length, residue composition and positional composition. Bars and whiskers, donor median and interquartile range. Each sampled model generated 500,000 sequences; exact BEAM marginals are also shown. BEAM and LZGraph [15] exclude Emerson. SoNIA (corpus) and SoNIA (shipped) [13] are corpus-refitted and default human TRB selection models; OLGA [11] is the default recombination model (Methods). (**h**) Length-15 logos: retained-count-weighted corpus composition (1.33 10^9^ observations, 192,275,347 distinct sequences) and BEAM’s exact positional marginals. Common information scale; colours, amino-acid chemistry.

**Figure 3:**
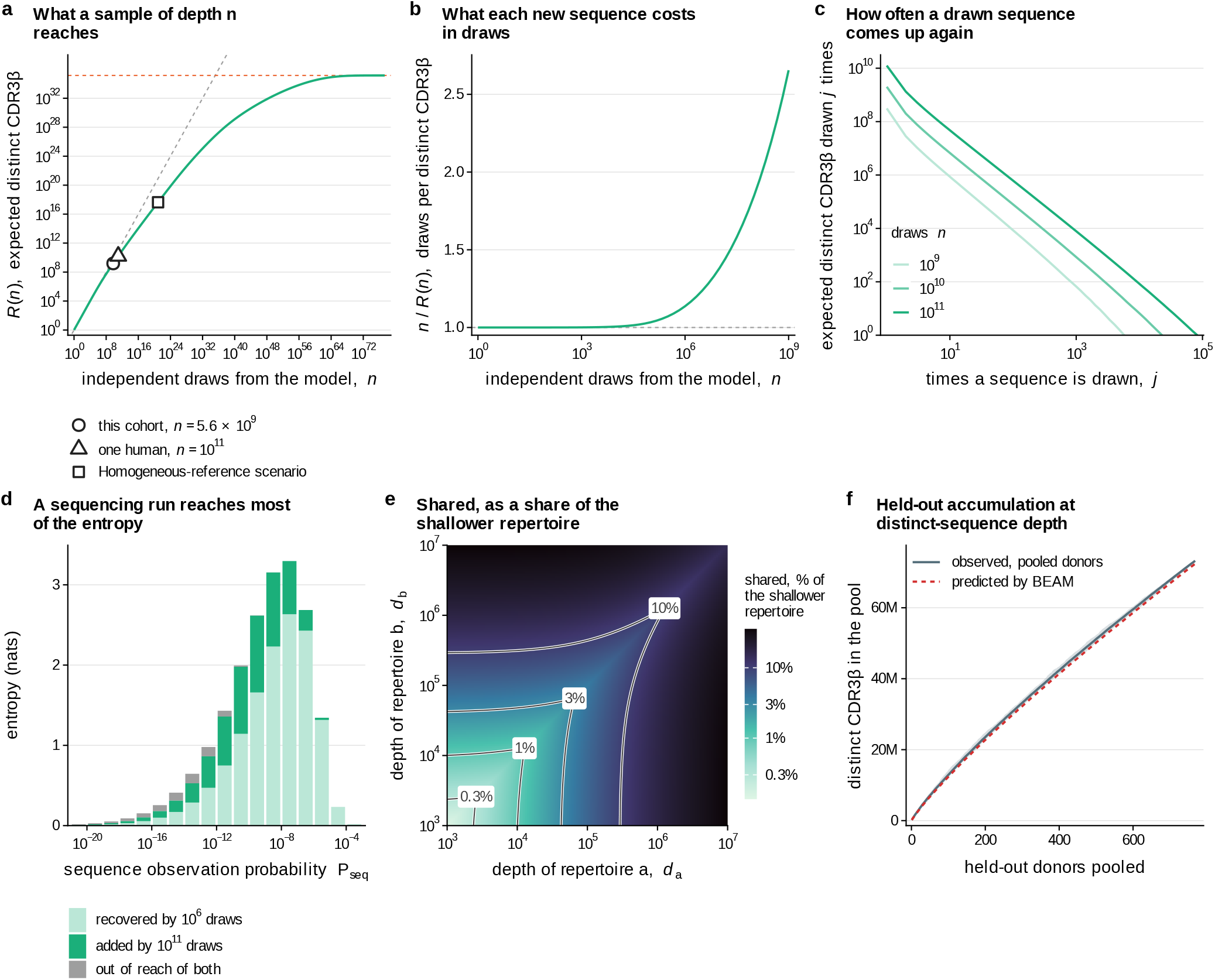
Finite sampling of the BEAM probability landscape. (**a**) Expected distinct strings R(n) after n independent draws from the population reference model (green; logarithmic axes). Dashed lines: R = n (grey) and model-domain size 1.41 *×* 10^35^ (orange). Reference depths: corpus (5,629,439,400 total input weight), adult naive-pool scale [19] (10^11^ draws) and a hypothetical homogeneous-reference scenario: 8.3 *×*10^9^ people *×*10^11^ independent draws per person from the same shared distribution (8.3 *×* 10^20^ draws in total). (**b**) Draws per expected distinct sequence n / R(n), to 10^9^ draws; dashed line, one; logarithmic x axis. (**c**) Expected sequences drawn exactly j times after 10^9^, 10^10^ and 10^11^ draws (light to dark), per unit j in logarithmic bins. The y axis starts at one expected sequence per unit j. This describes independent-sample multiplicity. (**d**) Model Shannon entropy by decade of sequence observation probability *P*_seq_: expected plug-in sample entropy at 10^6^ draws (light green), additional entropy recovered at 10^11^ draws (dark green) and remaining shortfall (grey). Full bar heights give each decade’s contribution to model entropy. (**e**) Expected shared sequences between independent samples of depths *d*_*a*_ and *db* (10^3^–10^7^ draws each), divided by the shallower sample’s expected distinct count. Colour, logarithmic; contours, 0.3%, 1%, 3% and 10%. This provides a statistical sharing baseline at specified depths. (**f** ) Pooled distinct CDR3*β* across k of 768 held-out Emerson donors [1]. Observed line and band: mean and range over 30 pooling orders. Dashed prediction: mean R at cumulative distinct-sequence depth over the same orders, using the Emerson-excluded model. Each donor’s distinct count defines its effective sampling depth here; a–e use independent draws. The corpus reference depth sums reported template counts where available and unit weights per distinct amino-acid sequence within each repertoire otherwise.

**Figure 4:**
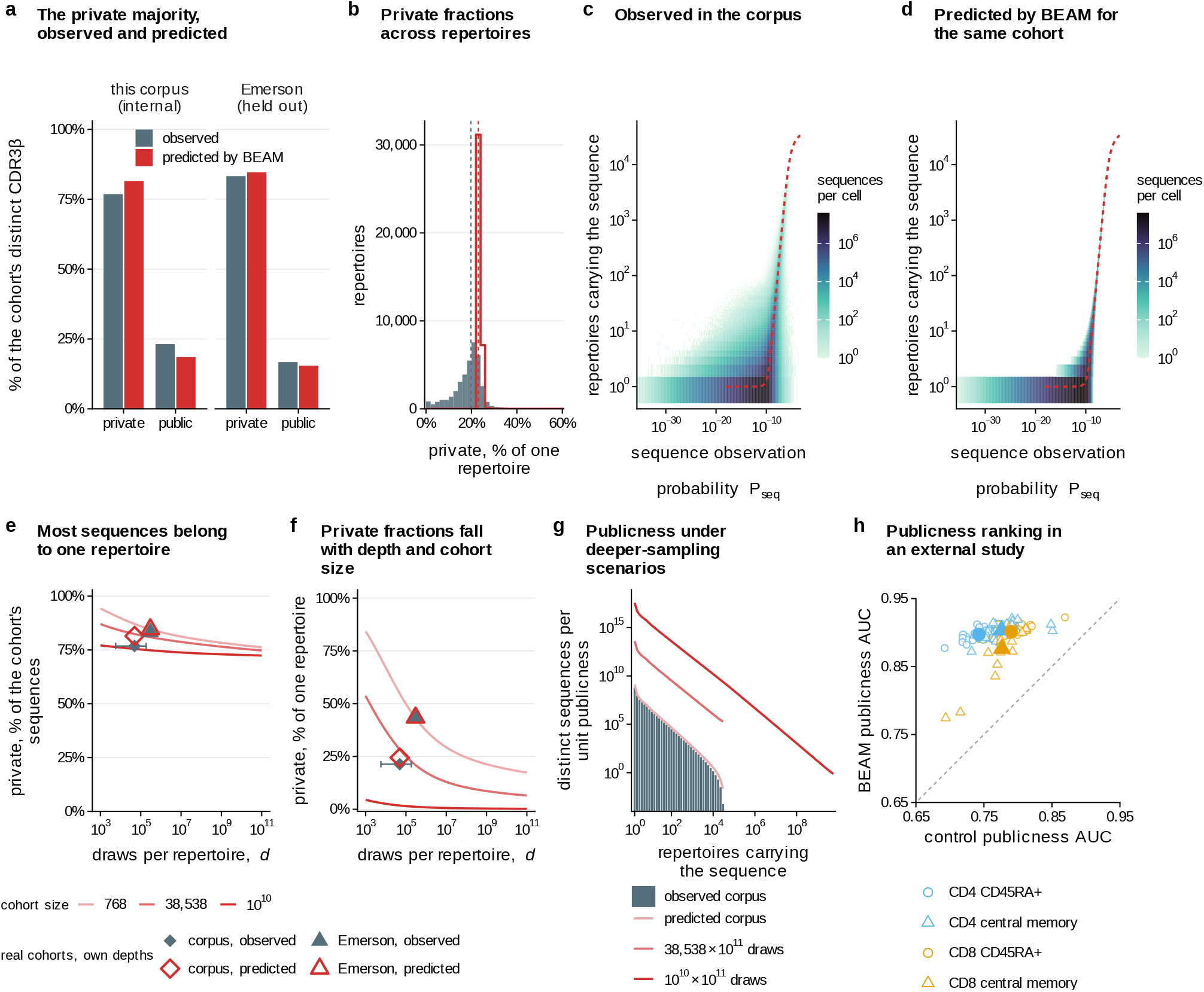
Observed and predicted sharing of CDR3*β* sequences. Private/public sequences occur in one/two or more repertoires of the specified cohort. Actual-cohort depths use reported template counts where available and unit weight per distinct amino-acid sequence within each repertoire otherwise. (**a**) Pooled private/public fractions: corpus (38,538 repertoires) and Emerson [1] (768 donors). Slate, observed; red, predicted. Corpus predictions are internal; Emerson donors were excluded from fitting. (**b**) Observed per-repertoire U/D (filled) versus E[U]/E[D] (outline), where U and D count private and distinct sequences. Dashed lines, medians; axis limit 60.0% contains 99.65% of observations. The ratios exclude sampling variation in U/D; held-out predictive intervals appear in Supplementary Fig. S4. (**c, d**) Observed (c) and predicted (d) sequence counts by *P*_seq_ and publicness, using 0.1-decade probability columns and a shared logarithmic colour scale. Panel d omits expectations below one. Dashed curve, mean publicness conditional on observation. (**e, f** ) Predicted private fractions versus equal repertoire depth for cohorts of 768, 38,538 and 10^10^ (light to dark): pooled private/pooled distinct (e); mean private/mean distinct within repertoires (f). Diamonds, corpus; triangles, Emerson; filled/open, observed/predicted at actual depths. Horizontal positions/bars show median/interquartile depths. The 768-repertoire curve and Emerson markers use the Emerson-excluded model; other curves use the population reference model. (**g**) Sequences per unit publicness in logarithmic bins: observed corpus (bars); predicted corpus and scenarios with 10^11^ draws per repertoire in 38,538 or 10^10^ repertoires (curves; legend gives repertoire count *×*draws per repertoire). Predictions use a discretised spectrum, with Poisson/normal approximations for the largest cohort; terminal model bins are omitted. Scenarios e–g assume independent draws from a shared fitted distribution; population-scale settings are homogeneous-reference scenarios. (**h**) External ranking by 4 naive/memory CD4/CD8 models trained on 10 Rubelt donors [18], tested in 30 ADC T1D donors [20, 21]. Catalogues contain 5,000 distinct sequences before excluding sequences seen in any training population. Public sequences recur in another donor’s same-fraction catalogue. Open symbols, donor/fraction ROC AUCs; filled, equal-donor means. Q is the same-fit independent-position control, preserving length and positional marginals. Dashed line, equal AUC. All 108 dependent comparisons favour BEAM. This assesses ranking, not sharing calibration or independence from V/J associations. Exact-length and catalogue-size controls appear in Supplementary Fig. S5.

**Figure 5:**
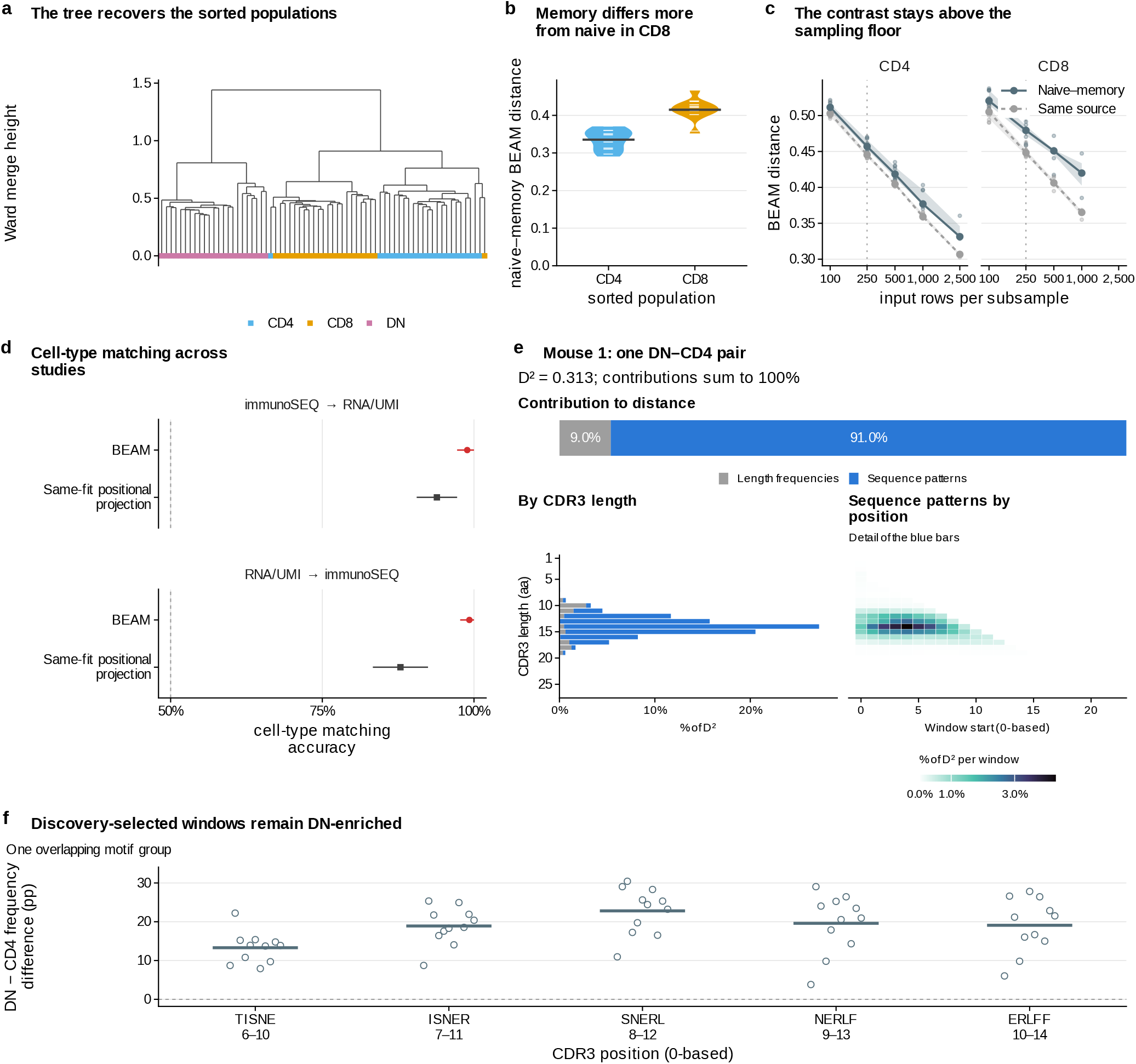
Repertoire comparison and interpretation with BEAM distance. Repertoire models use unit weight per clone record (a–c, e, f) or distinct CDR3*β* string (d). Distances compare length and local sequence distributions. (**a**) Ward dendrogram [27]: 69 repertoires from 23 mice [22, 17]; tip colours, CD4, CD8 or double-negative (DN). (**b**) Naive–memory distances in 10 donors from 5 twin families (reannotated Rubelt cohort [18]). Violins, donors; bars, means. (**c**) Naive–memory distances (solid) and disjoint same-repertoire subsamples (dashed) versus input depth. Lines/bands, donor medians/interquartile ranges; points, donors (3–10 per depth). Dotted line, primary depth of 250 clone records. (**d**) Cross-study CD4/CD8 nearest-neighbour matching at 1,000 distinct strings: 30 RNA/UMI [25] and 22 immunoSEQ donors [26] (Supplementary Fig. S11). Arrows, reference *→* query; red circles, BEAM; grey squares, its same-fit positional projection retaining length and single-position marginals. Points average donors after averaging CD4/CD8 queries and 3 subsamples within each donor. Whiskers, 95.0% query-donor bootstrap confidence intervals (10,000 draws; reference cohort fixed). (**e**) Mouse 1’s DN–CD4 distance (lowest ID; 174 DN and 213 CD4 records). Length (grey) and sequence-pattern (blue) contributions partition D^2^. Length bars sum to 100%; the heatmap expands blue contributions by five-residue windows (whole sequences below length five), with zero-based starts. (**f** ) Five overlapping windows selected by distance contribution in 12 discovery mice. Points, observed DN −CD4 window-frequency differences among length-15 clone records in 11 reserved mice; bars, unweighted mouse means; dashed line, zero. Labels give sequences and zero-based positions; horizontal offsets separate points. The windows form one group. Contributions, controls, alignment and post hoc complete-motif characterisation appear in Supplementary Fig. S12.

## 1 Results

### 1.1 A population reference sequence distribution

The corpus contains 926,909,993 distinct CDR3*β* amino-acid sequences from 38,538 repertoires, whose distinct-sequence counts span 4.5 decades (Fig. 2a, b; Supplementary Methods S1; Supplementary Table S1). Pooling all other repertoires covers a median 80.3% of a held-out repertoire’s sequences, with coverage still increasing; a median 19.7% occurs nowhere else (Fig. 2c). BEAM assigns observation probabilities to observed and unobserved sequences to calculate what samples of specified depths are expected to contain.

Positive probability extends to all 1–27-residue strings of standard amino acids, a mathematical domain of1.41 ×10 ^35^ strings. Yet sequences with *P*_seq_ ≥ 10 ^−12^ carry 91.7% of the probability while constituting only1.6 × 10 ^−25^ of this domain (Fig. 2d). Probability mass peaks at 15 residues, matching the abundance-weighted corpus, but probabilities span many decades within each length (Fig. 2f). In contrast, the20 ^*L*^ growth in possible strings places 95.0% of the mathematical domain at the rarely observed length of 27 residues.

A model built without the 768 Emerson donors [1] closely matches their length and residue marginals (Fig. 2g). BEAM gives the lowest median distance for sequence length and performs similarly to the corpus-refitted selection model [13, 14] (SoNIA (corpus)) for overall amino-acid composition; the latter is closer on positional composition. OLGA [11] describes generation before selection, a different target from the observational distributions evaluated here.

To test prediction beyond marginals, we fitted separate naive and memory CD4 and CD8 models while holding out both members of each of the 5 twin pairs in turn [18]. For sequences absent from every training donor, mean log probability exceeded an independent-position projection matching each fit’s length and positional marginals in all 40 donor–population comparisons (Fig. 2e). The gain, averaging donors, populations and families equally, was 3.66 nats per unseen sequence record and persisted after length normalisation and equal weighting of distinct strings (Supplementary Fig. S1a, b). Within-repertoire presence weighting and restriction to template-counted repertoires retained similar marginal agreement; bidirectionality improved unseen-sequence prediction over either direction alone, although three-residue contexts slightly outperformed four-residue contexts on this readout (Supplementary Fig. S2a, b).

Exact positional marginals recover the conserved ends and variable junction (Fig. 2h; Supplementary Methods S2). At length 15, mean positional total variation is 1.1% against the abundance-weighted corpus, versus 9.4% against distinct-sequence composition. The reference represents template-weighted observations where available and unit weight per distinct sequence within each repertoire otherwise (Methods 3.4).

### 1.2 Depth-dependent sampling expectations

BEAM computes *R*(*n*), the expected distinct-sequence count among *n* independent draws (Eq. (6); Fig. 3a; Supplementary Methods S4). At 10 ^11^ draws, a reference scale of the order of an adult’s naive T cell pool [19], it predicts 1.58 ×10^10^ distinct CDR3*β*. Approximately one in a hundred draws repeats a sequence by 2.2 ×10^4^ draws (Fig. 3b). At 10 ^11^ draws, 79.5% of distinct sequences appear once, while high-probability sequences can appear tens of thousands of times (Fig. 3c).

BEAM also quantifies the finite-sampling bias of plug-in Shannon entropy, which approaches the model’s entropy before distinct-sequence counts saturate. Expected plug-in entropy recovers 69.8% at 10 ^6^ draws (Fig. 3d; Supplementary Methods S5). Rare sequences dominate the unobserved count and contribute collectively to the entropy deficit. Agreement between direct calculations and the discretised spectrum is checked in Supplementary Fig. S3a, b.

The same baseline predicts overlap between samples (Eq. (8); Fig. 3e). Overlap, divided by the shallower sample’s expected distinct count, increases with depth: common sequences dominate small samples and are likely to occur in larger ones. Overlap reaches 35.4% for 10 ^3^ versus 10 ^7^ draws, compared with 0.14% for two samples at the smaller depth.

Without adjustment to the 768 held-out Emerson donors, BEAM predicts 98.9% of their observed pooled distinct-sequence count (Fig. 3f). Depth here sums each donor’s distinct amino-acid sequence count; the publicness analysis below uses abundance-based depths (Methods 3.1). Globally counting each corpus sequence once overpredicted accumulation; the primary configuration was closest at the final endpoint among those tested (Supplementary Fig. S2d).

### 1.3 Observation probability and publicness

Publicness counts the repertoires carrying a sequence within a specified cohort; sequences found in exactly one are private. Most pooled distinct CDR3*β* are private: 76.8% in the corpus and 83.3% among Emerson donors (Fig. 4a). Using template counts where available and unit weight per distinct sequence within each repertoire otherwise, BEAM predicts 81.5% and 84.6%, respectively, without fitting to publicness; the Emerson prediction uses the excluded-cohort model. Under these abundance-based depths, predicted pooled distinct-sequence counts are 1.68 and 1.50 times the observed counts in the corpus and Emerson cohort, respectively.

Across corpus repertoires, the observed private-fraction interquartile range is 6.5 times that of the depth-specific ratios of expected private to expected distinct counts (Fig. 4b). In complete-cohort simulations of the separate held-out Emerson cohort at each donor’s original draw count, the 95% predictive interval for the across-donor interquartile range was 0.99–1.08 percentage points, compared with 2.23 observed (Supplementary Fig. S4a, b). Observed donor variation thus exceeded that reproduced by sampling from the fixed shared distribution.

BEAM reproduces the broad increase in publicness with observation probability (Fig. 4c, d). Publicness ranking persisted across model-design and weighting changes in sampled Emerson catalogues, including training-seen strings (Supplementary Fig. S2c).

To examine the effects of depth and cohort size, we calculated publicness under independent sampling from a shared fitted distribution. Across the evaluated scenarios, private sequences remain a majority of the pooled catalogue, while their fraction within each repertoire falls as the cohort grows (Fig. 4e, f). Numerous rare sequences occur in only one repertoire, whereas frequently sampled sequences recur across repertoires. The predicted publicness spectra decline approximately as power laws over the fitted range 10–3,162. Deeper sampling increases expected counts across the spectrum, while larger cohorts extend its publicness range (Fig. 4g; Supplementary Methods S4).

For an external test, we fitted population-specific models to 10 Rubelt donors [18] and ranked public versus private sequences in 30 donors from an independent AIRR Data Commons dataset [20, 21]. Among strings absent from every training population, the mean area under the receiver operating characteristic curve ranged from 0.88 to 0.90 across sorted fractions; the corresponding independent-position projections ranged from 0.74 to 0.79. All 108 donor–fraction comparisons favoured BEAM over its independent-position projection (Fig. 4h). The advantage persisted within exact lengths and in smaller catalogues (Supplementary Fig. S5a; Methods 3.8).

Probability-weighted sequence logos show increasingly constrained composition with expected publicness, first at the N-terminal end and then through the junction (Supplementary Fig. S6a).

### 1.4 Repertoire comparison from sequence distributions

We fitted one BEAM model per repertoire and compared length distributions and local block marginals of up to five residues by Hellinger distance (Eq. (10); Supplementary Methods S6). The distance includes patterns absent from either sample and decomposes into contributions from length and local composition.

Clustering 69 mouse repertoires [22, 17] places 67 with their sorted CD4, CD8 or double-negative (DN) population (Fig. 5a), matching TCRdist with optimal transport, which also uses V genes and pairwise sequence similarity [16, 17] (Supplementary Fig. S7a).

In all 10 donors from 5 twin families [18], naive–memory distance is larger in CD8 than CD4 (Fig. 5b), matching the Repertoire Dissimilarity Index [5] (Supplementary Fig. S7b). The family-level two-sided sign-flip *P*=0.0625 is the minimum attainable with five families. This ordering persisted across model-design and weighting changes (Supplementary Fig. S2e). At the predeclared 250 clone records, naive–memory distances exceed distances between disjoint samples from the same repertoire by 0.0132 in CD4 (0.00441–0.0220) and 0.0293 in CD8 (0.0229–0.0357; 97.5% family-jackknife confidence intervals; Fig. 5c; Methods 3.9). Separation also persists at unequal depths (Supplementary Fig. S8a, b).

Controlled immuneSIM perturbations [23] support sensitivity to junctional insertions, D-gene trimming and implanted motifs; V/D/J usage noise did not significantly separate conditions in this simulation (Supplementary Fig. S9a). In a descriptive longitudinal example of two donors [24], all 72 repertoires’ nearest neighbours came from the same donor. For peripheral blood mononuclear cell repertoires, temporal changes were smaller than between-donor differences (Supplementary Fig. S10).

Larger human cohorts comprised paired CD4/CD8 repertoires from 30 people in an external RNA study using unique molecular identifiers (RNA/UMI) [25] and 22 people in an immunoSEQ study [26]. Models used equal weight per distinct amino-acid sequence. At 1,000 sequences, every donor’s mean distance to others was smaller within than between cell types (Supplementary Fig. S11a). Using references entirely from the other study, nearest-neighbour accuracy was 98.9% for RNA/UMI queries and 99.2% for immunoSEQ queries, versus 93.9% and 87.9% for the same fit’s positional projection (Fig. 5d). This control retains length and single-position marginals but omits local residue combinations. Accuracy remained high across 1,000–5,000 distinct sequences in fixed donor subsets (Supplementary Fig. S11b–d). Gene-usage methods were stronger in some within-study comparisons, and empirical positional composition was competitive (Supplementary Methods S7).

In the illustrated mouse DN–CD4 pair, 91.0% of squared distance comes from patterns within lengths, with the remainder from length frequencies (Fig. 5e). Ranking window contributions across 12 discovery mice selected five overlapping five-residue windows at length 15. Each was DN-enriched in observed within-length frequencies in all 11 reserved mice (Fig. 5f). These correlated windows form one overlapping group; their contributions, controls and post hoc complete-motif characterisation are detailed in Supplementary Fig. S12 and Supplementary Methods S6.

## 2 Discussion

BEAM connects sequence observation probabilities to sampling expectations and interpretable repertoire comparison. A population reference model predicts distinct-sequence counts, overlap and publicness at specified depths; separate models fitted to individual repertoires trace differences to lengths and local patterns. Both uses describe observational distributions, connecting what finite samples contain with how repertoires differ.

The local, bidirectional structure balances predictive information with tractable computation. The gain in held-out sequence prediction over a control with identical length and individual-position distributions shows that learned residue relationships carry useful information beyond those summaries. V(D)J generation and selection models connect sequence probabilities to biological mechanisms [9, 13, 14]. BEAM complements these approaches by fitting the observational distribution directly from sequence counts, without requiring gene annotations. The appropriate model order depends on the task and computational budget.

With a population reference, researchers can examine how additional sampling changes expected sequence recovery and overlap. Rare sequences can dominate the pooled catalogue even when individual repertoires contain many sequences shared with others, making publicness dependent on both depth and cohort size [4]. Held-out accumulation and external publicness ranking support these uses of fitted reference distributions under the evaluated input conventions. Abundance-based count overprediction and donor variation beyond the predictive sampling intervals identify departures from the shared reference (Supplementary Fig. S4). Donor-specific expansion, repertoire heterogeneity and measurement conventions are possible contributors whose individual effects remain unresolved. A researcher could, for example, fit BEAM to a template-counted pilot repertoire and calculate expected sequence recovery and replicate overlap at candidate sampling depths. These calculations can identify where additional sampling has diminishing returns, while predictive intervals from subsequent samples can assess compatibility with the fitted sampling model.

The same fitted distributions support repertoire comparison through length and local-block marginals. Cross-study CD4/CD8 matching supports transfer across the evaluated human datasets. The sources differ jointly in assay, processing and clinical cohort, so the benchmark does not isolate their individual effects (Supplementary Fig. S11). Decomposing squared distance then provides a path from repertoire separation to specific sequence features. The mouse example shows reproducible DN enrichment of discovery-selected windows belonging to one overlapping group (Fig. 5f). Such attribution can prioritise correlated sequence features for further biological investigation.

The model’s scope follows from its inputs and assumptions. The reference inherits the populations and measurement conventions represented in the corpus. Combining template counts with sequence presence changes the meaning of a model draw. Observation probabilities reflect recombination and selection together with expansion retained through abundance weights; recurrence can also arise from convergent recombination [7, 8]. Sampling extrapolations assume independent draws from the specified distribution. The current model describes CDR3*β* without explicit paired-chain or human leukocyte antigen (HLA) context, and its finite local structure does not explicitly represent arbitrary long-range interactions. Its mathematical sequence domain includes strings that need not be functional receptors. Computational exactness is consequently distinct from biological calibration, and extreme-depth projections remain conditional scenarios.

Reference models stratified by cell population or host characteristics, together with harmonised abundance measurements, could test whether the residual publicness variation can be reduced. Future work could extend and validate the framework for B cell receptor heavy and light chains, other TCR chains and nucleotide-level repertoire analysis. Paired-chain and longer-context models offer further directions, with predictive gains assessed alongside computational cost. The C implementation and Python interface make the present calculations reusable in repertoire analyses, with resource requirements documented in Supplementary Table S2. BEAM makes sampling-dependent properties of fitted TCR sequence landscapes directly computable and connects repertoire-level differences to local sequence features that can be tested experimentally.

## 3 Methods

### 3.1 Data

#### Corpus

We aggregated 38,538 human TCR*β* repertoires from 447 published immunosequencing studies spanning 2009 to 2025, drawn from six public sources (Supplementary Table S1): the Adaptive Biotechnologies immuneACCESS database [28] (immunoSEQ assay;73.7%of repertoires), TCRdb 2.0 [29] (16.4%), the AIRR Data Commons [20] (7.1%), mal-ID [30] (1.4%), VDJbase [31] (1.2%) and Britanova et al. [32] (0.2%). The corpus is a literature aggregate rather than a healthy reference cohort.

Repertoires were retained if they were human, held at least 100 sequence records after the sequence filters, and passed the duplicate-content screen. Because public repositories re-host one another extensively, duplicate candidates were identified by comparing the CDR3*β* sets of all 1,317,923,470 repertoire pairs rather than by metadata, which removed 94 exact and 11,606 near-identical repertoires (Supplementary Methods S1).

Within each repertoire we kept CDR3*β* amino-acid sequences of 3 to 27 residues spelled entirely from the twenty standard amino acids, and merged the records of each amino-acid sequence. This is a stop-free alphabet filter, rather than a test of rearrangement productivity: records annotated as out of frame are kept if their amino-acid CDR3 is spelled from standard residues, and no filter is applied to the conserved cysteine or the terminal phenylalanine or tryptophan, which 99.6% and 98.6%of sequences carry. The alphabet test removes 50,262,483 distinct sequences (5.1%) containing stop codons, ambiguity codes or gap characters, and the upper length bound follows the measured length distribution, whose main mode ends sharply at 27 residues (Supplementary Methods S1). The corpus holds 926,909,993 distinct CDR3*β* amino-acid sequences.

#### Abundance

The repositories report abundance in different units: platform-reported estimates of input-template counts for immunoSEQ repertoires [33, 34], and sequencing reads or record counts elsewhere, which need not be proportional to input molecules. We therefore kept template counts where the platform reported them (27,965 repertoires) and assigned unit weight per distinct aminoacid sequence within each of the 10,573 repertoires without them. This minimum-count convention gives a total observation weight of 5,629,439,400;69.3 % of the corpus-wide distinct sequences have total weight one, against a mean of 6.1 per sequence. For sampling and publicness, depth is the sum of these abundance weights, and one unit is one draw from the model. For repertoires without template counts, this depth is the number of distinct sequences, not an estimate of the number of input molecules. Two other units are used explicitly: distinct amino-acid sequences describe repertoire size in Fig. 2b and define the input depth of the held-out accumulation in Fig. 3f; the mouse and naive–memory repertoire-distance benchmarks use clone records, with one unit per record. The larger human CD4/CD8 benchmark (Fig. 5d; Supplementary Fig. S11) instead weights each distinct amino-acid sequence once. These quantities are kept separate because multiple records or templates can encode the same amino-acid sequence.

#### Held-out cohort

The 768 donors of Emerson et al. [1] in the corpus served as a held-out cohort. A second model was built without them (§3.4) and evaluated on them (Figs. 2g, 3f and 4a, e, f; Supplementary Fig. S4). Fig. 4b instead compares the full corpus with the population reference model.

#### Repertoire-distance benchmarks

The mouse, naive–memory, simulation and longitudinal comparisons used four datasets with known structure:69TCR*β* repertoires of 23 mice, each sorted into CD4, CD8 and DN T cells (DN being the intestinal CD4^−^CD8*αβ*^−^CD8*αα*^+^ subset), generated by Schattgen et al. [22] and tabulated by Olson et al. [17]; the naive and memory CD4 and CD8 TCR*β* repertoires of 10 donors in five twin families [18], using a public reannotation of the cohort analysed with the Repertoire Dissimilarity Index [5]; new repertoires simulated with immuneSIM 0.8.7 [23], adapting the controlled-perturbation benchmark of immuneREF [35]; and72 TCR*β* repertoires of two donors sampled over time, from the published archive of Minervina et al. [24], represented by selected top-clone records (611 to 10,000 records per repertoire).

The larger human CD4/CD8 benchmark (Fig. 5d; Supplementary Fig. S11) used paired blood repertoires from 30 people in GSE285951 [25] and 22 people with multiple sclerosis from Emerson et al. [26]. The former was newly downloaded from GEO; its accession and sample aliases were absent from the checked corpus registry. The latter was already represented in the corpus. Models were fitted separately to each repertoire, without using a corpus model. Source filtering, pairing and sampling are detailed in Supplementary Methods S7.

### 3.2 The BEAM model

#### Notation

A sequence in the model domain is a 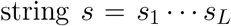 of length *L*=|*s*| over the alphabet *A* of the 20 standard amino acids, with 1≤ *L* ≤ *L*_max_ = 27. We write*s* 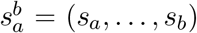 for a substring and set*s i* =*∅*, a boundary symbol, for *i* <1and*i* > *L*, so that substrings near either end are padded with *∅*.log denotes the natural logarithm and log_10_ the base-10 logarithm; one decade corresponds to a tenfold change in probability.

#### The model

BEAM scores every residue twice, by two experts. Each expert is a table of distributions over *A*, with one row for every position *t*= 1, …, *L*_max_, counted from the expert’s own end of the CDR3, and every context *u* of *k* neighbouring symbols. The forward expert *q*_*F*_ (*a* | *t, u*) reads from the start: for residue *si* it uses *t*=*I* and the *k* symbols before it, 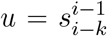. The backward expert reads *q*_*F*_ (*a* | *t, u*) reads from the end: it uses *t* = *L* − *i* + 1 and the *k* symbols after it, 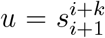 Neither table depends on *L*, so one set of parameters serves every length, as the left and right features of SoNIA do [13]. The sequence observation probability of *s* is

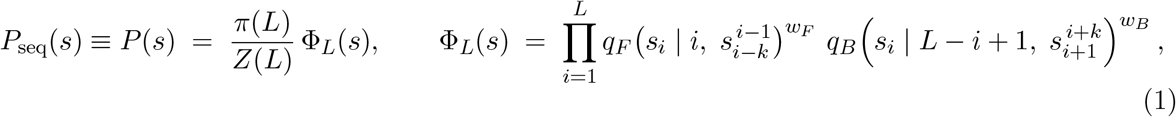

where *π*(*L*) is the distribution of lengths and

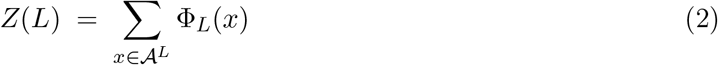

sums the weight Φ*L* over all 20^*L*^ strings of length *L*. Each expert is normalised row by row, but their product is not: Φ _*L*_ does not sum to one over *A* ^*L*^, and dividing by *Z*(*L*) makes *P* (· |*L*) = Φ _*L*_*/Z*(*L*) a distribution over the strings of length *L* . Hence ∑ _|*s*| = *L*_ *P*(*s*) = *π*(*L*) and ∑ _*s*_ *P*(*s*) = 1. We use order *k* = 4 and weight *wf* = *wB* = 1/2, so that each residue is scored by the geometric mean of the two experts’ probabilities, a product of experts [36]. We retained order four because it provides the longest context supported by the implementation and enables five-residue feature attribution, while giving broadly comparable performance across held-out fidelity, accumulation, publicness and repertoire-distance analyses; order three performed slightly better on the specific unseen-sequence cross-entropy readout; equal weights give the two sequence directions equal influence. These settings are fixed across the primary analyses; sensitivity to order, direction and corpus weighting is evaluated in Supplementary Fig. S2 and detailed in Supplementary Methods S8. The two experts parameterise one distribution; they are not models of separate biological processes.

#### Exact normalisation

Each factor of Φ_*L*_ involves at most *k*+1 consecutive symbols. *Z*(*L*)is therefore a sum over the paths of a layered graph whose nodes are the last *k* symbols written, 21^*k*^ nodes per layer counting *∅*, and a forward dynamic program computes it exactly in *O*( *L* ·21·^*k*^ 20) operations rather than 20^*L*^ (Supplementary Methods S2). The same graph, swept forward and backward, gives exact marginal distributions of residues at any set of consecutive positions. No sequence-space sampling or pruning is required. We use *P* (*s*) as shorthand for *P*_seq_ (*s*) in the derivations. This is the normalised probability of *s* on one draw from the fitted observation distribution; its probability of appearing at least once in *n* independent draws is 1− (1 −*P*(*s*))^*n*^. Here, exact refers to the complete finite sum, evaluated in floating-point arithmetic.

### 3.3 Expert tables and hierarchical prior

#### Counts

A model is built from input sequences *s* with weights *ω*_*s*_: the mixed template/presence weights of the population reference model, or the record or distinct-sequence weights specified for each repertoire benchmark (§3.4). Each input sequence contributes its weight once per residue to each expert. For the forward expert, the count of residue *a* in row(*t, u*) is

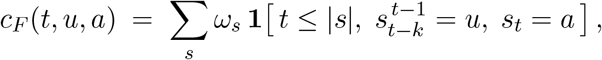

and *c*_*B*_ (*t, u, a*)is the same sum with the residue *s*_*i*_,*I* = |*s*| −*t*+ 1, in place of *s*_*t*_ and its context 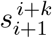 in place of 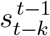. The tables have *L*_max_ ·21^*k*^ rows per expert, most of them observed rarely or never.

#### Hierarchical prior

Each row is estimated by hierarchical Dirichlet backoff [37], which shrinks it towards the row with the same position and a shorter context. Let *u*_*j*_ be the *j* symbols of *u* nearest to the scored residue (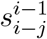 for the forward expert, 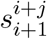 for the backward), and let *c*(*t, u*_*j*_, *a*)be the counts summed over the *k* − *j* symbols that *uj* drops, with *c*(*t, u*_*j*_) = ∑*a c*(*t, uj, a*). Separately for each expert and for *j* = 0, …, *k*,

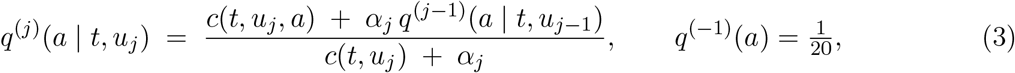

and the expert is the top level, *q*(*a*| *t, u*) =*q* ^(*k*)^(*a* |*t, u*_*k*_). Level 0 conditions on position alone. The strength *α*_*j*_ >0 is the number of observations that the shorter context is worth at level *j* : a row with many observations follows its own counts, a row with few leans on its parent, and a row never observed equals its parent. Both experts use the same strengths. The length distribution is estimated in the same way against a uniform parent,

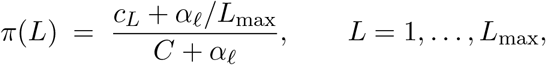

with *c*_*L*_ the weight of input sequences of length *L* and *C* =∑_*L*_ *cL*. The expert tables follow from one pass of counting and the closed-form estimates of Eq. (3); each prior strength is selected by a one-dimensional evidence search, described below.

#### Estimating the strengths

The strengths are selected by an empirical-Bayes rowwise evidence score. If a row’s residue distribution has a Dirichlet prior with mean equal to the parent’s prediction and concentration *α*_*j*_, Eq. (3) is its posterior mean. Integrating out the row’s distribution while holding the estimated parent fixed gives its Dirichlet–multinomial evidence; summing over rows at level *j* yields

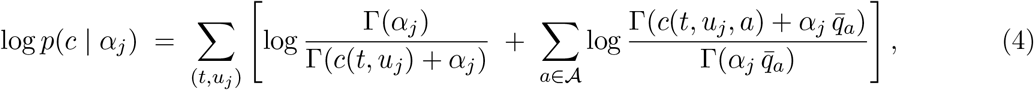

where 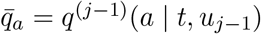 is the parent’s prediction and the sum runs over the observed rows of level *j* in both experts, up to a term that does not depend on*α*_*j*_ (Supplementary Methods S3). This conditional score assumes independence across rows and observations. The levels are estimated in order, *j*= 0, …, *k*, each against parents built with the strengths already estimated. The search is performed on the logarithmic strength, maximising Eq. (4) plus the Gaussian log density of a weak hyperprior, log *α*_*j*_*∼ N* (log 10,(log 100) ^2^), which is most influential when the counts provide little information about the strength; *α*_*ℓ*_ is estimated in the same way.

#### Support

Every row is blended, level by level, down to the uniform distribution, and *π*(*L*) > 0 for every *L*. Every table entry is therefore positive, every string of 1 to 27 residues has *P* (*s*) > 0, and the support of the model is all 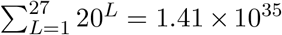 strings. This is a mathematical domain for smoothing and computation, not a count of biologically realisable receptors. The corpus filters retain lengths 3–27; the positive prior extends model support to lengths 1 and 2.

### 3.4 The models

The population reference model was built from the whole corpus, with each distinct sequence weighted by its summed template/presence weight (§3.1). It defines the corpus-wide probability landscape and provides the reference for sampling and publicness calculations (Fig. 1c). Its estimated strengths are (*α*_0_, …, *α*_4_) = (5.19, 9.77, 24.99, 83.05, 77.99) and *α*_*ℓ*_ = 4.14: the fitted prior strengths are larger for high-order contexts than for position-only rows. The held-out model was built in the same way after the Emerson donors’ abundances were subtracted from the corpus, which removes the 32,520,669 sequences found only in them and lowers the abundance of 40,725,007 others. It was built from 894,389,324 distinct sequences, with strengths within 2% of the population reference model’s, and is used wherever predictions are compared with that cohort. For repertoire distances (Fig. 5), one model was built from each repertoire with the same order, length range and prior specification. Clone records received equal weight in panels a–c, e and f; each distinct aminoacid string received unit weight in panel d. The weighting defines what a model describes: the population reference combines reported template abundance and sequence presence, whereas the Fig. 5 models describe repertoire sequence composition at the stated input unit. The population reference model is consequently an observational population model that can reflect recombination, selection, exposure and expansion. Independent sampling from it is a statistical reference and does not separate those biological processes.

### 3.5 Sums over all sequences

Sampling expectations and probability-spectrum summaries are sums of functions of *P* (*s*) over the model domain of 1.41 × 10^35^ sequences. These whole-space sums are evaluated without drawing sequences from the model, using the two computations below. Separate simulations assess marginal fidelity and predictive sampling intervals.

#### The probability spectrum

For a set *B* of values of log *P*, let *N*_*L/*_ (*B*) be the number of sequences of length *L* with log *P*(*s*) ∈ *B* and *P*_*L*_ (*B*) the probability they carry. Because log Φ _*L*_ (*s*) is a sum of terms along the sequence, the forward dynamic program can carry at each node a histogram of the accumulated log Φ*L* on a fixed grid instead of a single sum. One pass then gives discretised *N*_*L*_ and *P*_*L*_, preserving the total sequence count and probability mass. On the 32,768-point grid used for the full model, each sequence is placed within 0.09 decades of its exact log_10_ *P*. Sums of the form ∑*N f*(*P*(*s*)) are then evaluated as ∑ *N* (*B*) *f* (*pB*) over the non-empty bins, with *N* (*B*) = ∑_*L*_ *N*_*L*_(*B*) and *p*_*B*_ the probability at the bin. The spectrum gives Fig. 2d, f and the quantities of Figs. 3c, d and 4c, d, g that depend on ranges of *P*_seq_. Banded forward and backward sweeps likewise give numerical estimates of positional composition within probability ranges (Supplementary Fig. S6a); band boundaries depend on the grid, unlike unconditional positional marginals.

#### Exact occupancy

A sum such as the expected number of distinct sequences in a sample, ∑ _*s*_[1− (1 −*P*(*s*)) ^*n*^], cannot be computed by the dynamic program directly, because 1 − (1 −*P*(*s*))^*n*^ does not factor along the sequence. Powers of *P*(*s*) do: *P*(*s*) ^*z*^ =*π*(*L*) ^*z*^*Z*(*L*)^−*z*^Φ _*L*_ (*s*)^*z*^, and Φ _*L*_ (*s*) – ^*z*^ is the product of the same factors each raised to the power *z*, so the power sum *M* (*z*) =∑_*s*_ *P*(*s*) ^*z*^ is computed exactly by the same dynamic program at any real or complex *z*. Each occupancy quantity is obtained from *M*: by a binomial series in *M* (1), *M*(2), …at small depths, and otherwise by a Mellin–Barnes contour integral, which expresses it as an integral of *M* along a vertical line in the complex plane. Numerical evaluations carry recorded error bounds (Supplementary Methods S4). These evaluations avoid spectrum discretisation and give expected distinct-sequence counts, sharing and private fractions (Figs. 3a, b, e, f and 4a, b, e, f) and check the values computed on the spectrum. For the population reference model, comparisons with the counting spectrum are shown for distinct-sequence counts (Supplementary Fig. S3a) and singleton fractions, model entropy and reconstructed probability mass (Supplementary Fig. S3b).

### 3.6 Model description and fidelity

#### Corpus and coverage

(Fig. 2a–c). Lengths are counted over distinct sequences (Fig. 2a), and repertoire size is its number of distinct amino-acid sequences (Fig. 2b). For coverage, we held out 300 repertoires and pooled the remaining 38,238 in 10 random orders. For each held-out repertoire and pool size we recorded the share of its distinct sequences already present in the pool, and the share found in no other repertoire of the corpus, which pooling cannot reach.

#### Fidelity to held-out repertoires

(Fig. 2g). Each model generated 500,000 sequences. For each of the 768 held-out donors we compared the generated and observed distributions of CDR3 length, of amino-acid composition, and of amino-acid composition at each of the first 18 positions counted from the start (pooled over lengths), by the Jensen– Shannon distance between distributions *µ* and 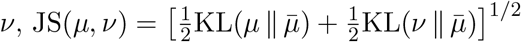with 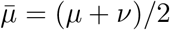, averaging the positional distances over positions with support in both distributions. Observed distributions count each in-frame source record of the donor once, retaining repeated amino-acid sequences. The models were the held-out BEAM model; OLGA with its default human TRB model [11], which describes generation before selection; the shipped and corpus-refitted human TRB SoNIA selection models, both using the `Sonia` class in soNNia 0.4.0 [13, 14]; and an amino-acid positional LZGraph [15] built from the corpus without the Emerson donors. The shipped and refitted SoNIA models were initialised with `ppost_model=humanTRB` and `pgen_model=humanTRB`, respectively. For the refit, 1,000,000 candidate CDR3/V/J records with resolvable gene annotations were sampled from a randomly selected pool of 1,000 non-Emerson repertoires, with a cap of 1,500 records per repertoire and one record per distinct CDR3 amino-acid sequence. Sampling stopped at the target size. The package’s default human TRB gene and sequence filters retained 946,490 records. The refit used the default additive features for length, joint V/J usage and residue identity at positions from each CDR3 end (maximum positional depth 25, maximum length 30). Training used up to 50 epochs, batch size 5,000, a 20%validation split, early-stopping patience 30, `set_gauge=True`, seed 20260831 and 1,000,000 generated background sequences, of which 999,996 passed filtering. Sequence generation was not conditioned on specific V or J genes. BEAM’s three distributions were also computed by exact marginalisation, which is the limit of infinitely many generated sequences. These readouts assess length and residue marginals rather than the full joint sequence distribution.

For the separate family-held-out sequence test (Fig. 2e), we used the processed Rubelt clone records described in §3.9. Each of 5 twin pairs was excluded in turn. For each of 4 populations—naive and memory CD4 and CD8—one model was fitted to pooled records from the other 8 donors, giving 20 population-specific models. Each nucleotide-defined clone key received unit weight, retaining identical amino-acid strings from different keys; molecular abundances were unavailable. Order four, equal directional weights, maximum length 27 and training-only evidence estimation of the prior strengths were fixed across fits. For a fitted distribution 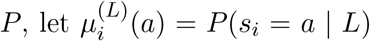. Its independent-position projection was

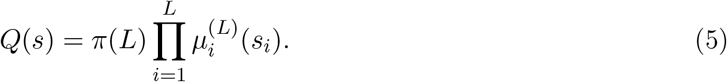

The marginals were computed exactly by dynamic programming. Thus *Q* is normalised and shares *P*’s length distribution and every individual-position marginal; it is an independent-position control constructed from the same fit. We scored each held-out record by log *P* (*s*) −log *Q* (*s*). The primary set excluded amino-acid strings observed in any population of the training donors. Scores were averaged within donor, then equally across twins, populations and families; the figure displays family means within each population. Scores divided by sequence length and equal weight per distinct amino-acid string within each test repertoire provide the controls in Supplementary Fig. S1a and b, respectively. The complete held-out set was also retained as a sensitivity summary. Training folds overlap and populations within a donor are dependent; no inferential tests were added.

#### Positional composition

(Fig. 2h). At length 15, BEAM’s exact positional marginals *P* (*s*_*i*_ =*a* |*L*)were compared with the abundance-weighted residue frequencies of the corpus at each position, by the total variation distance 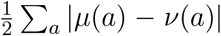 between the two compositions *µ* and *ν*. Logo heights scale each position’s composition by its information, log_2_ 20 bits minus the entropy of the composition in bits.

### 3.7 Sampling

A sample of depth *n* is *n* independent draws from the fitted observation distribution *P*. Independence is imposed on the draws; it does not imply that the data used to estimate *P* arose from independent recombination events. Writing *p* for *P* (*s*)inside the sums, the expected numbers of distinct sequences, of sequences drawn exactly *j* times, and of sequences found in both of two independent samples of depths *n*_*a*_ and *n*_*b*_ are

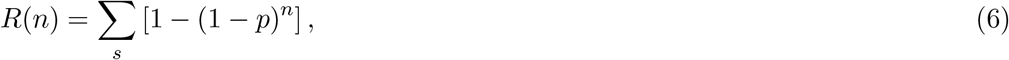

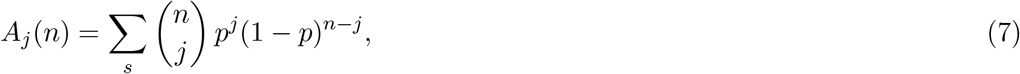

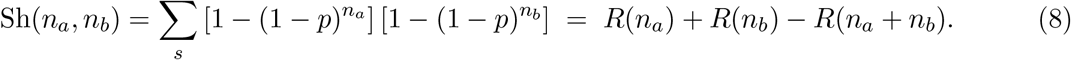

Fig. 3b shows the draws per expected distinct sequence, *n/R*(*n*), and Fig. 3e the shared sequences as a share of the shallower sample, Sh*/*min *R*(*n*_*a*_), *R*(*n*_*b*_) . The *A*_*j*_ spectra use Poisson occupancy. On the same spectrum, the Poisson and binomial evaluations of the total distinct-sequence count differ by less than 4 ×10^−8^ (relative) across the checked depths. Total-count and singleton summaries were evaluated directly from power sums. The entropy of the model, *H*=∑ _*s*_ *P*(*s*) log *P* (*s*) in nats, is computed exactly by the dynamic program. Under the Poisson approximation, expected plug-in entropy at depth *n* falls short of *H* by ∑_*s*_ *p δ*(*np*), where *δ*(*λ*) =E[log(1 +*X*)] – log *λ* for *X ∼* Poisson (*λ*) (Fig. 3d; Supplementary Methods S5). The 10^11^-draw reference is of the order used for one adult’s naive T cell pool [19]. The additional marker in Fig. 3a, labelled “Homogeneous-reference scenario”, illustrates the calculation at a larger pooled depth under independent draws from the same fitted distribution; its assumed cohort size and depth are specified in the caption. These are reference sampling scales, rather than measured numbers of independently generated receptors.

#### Held-out accumulation

(Fig. 3f). The 768 held-out donors were pooled one at a time in 30 random orders, recording the observed number of distinct sequences in the pool. The prediction for each order is *R* of the held-out model at the pool’s cumulative depth, averaged over orders. For this presence-based comparison, a donor’s input depth is its number of distinct amino-acid sequences, rather than its template count. This counts each observed amino-acid identity once per donor and limits the influence of within-donor abundance on the input depth. It is not a count of recombination events: different rearrangements can yield the same amino-acid sequence. Sampling and publicness calculations elsewhere use the abundance-based depths defined above.

### 3.8 Publicness

The publicness *m* (*s*) of a sequence is the number of repertoires in a cohort that contain it; a sequence is private relative to that cohort if *m*(*s*) = 1. A repertoire is an assay sample and is not necessarily an independent individual; donor identity is explicit for the Emerson cohort. Observed publicness was counted with exact sequence keys over the per-repertoire files of the corpus.

For prediction, a cohort is *N* repertoires with depths *n*_1_, …, *nN* and total *T*= ∑_*r*_ *nr*, each an independent sample from *P*. A sequence of probability *p* is present in repertoire *r* with probability

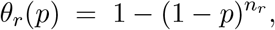

independently across repertoires. The publicness *m*(*s*)is therefore the sum of *N* independent Bernoulli variables with these probabilities, a Poisson-binomial variable. The expected numbers of distinct and of private sequences in the cohort are

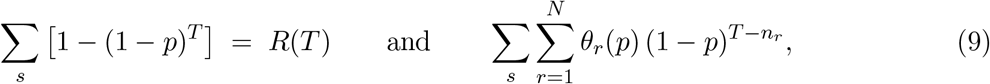

and the private count divided by the distinct count defines the predicted cohort private fraction (Fig. 4a). The corresponding ratio for repertoire *r* is 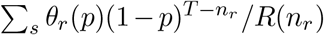 (Fig. 4b). The two fractions have different denominators, all distinct sequences of the cohort and the distinct sequences of one repertoire, and differ greatly. These predictions are ratios of expected counts, not expectations of the corresponding random ratios. Equal-depth scenarios and publicness spectra (Fig. 4e–g) are described in Supplementary Methods S4.

For the predictive analysis (Supplementary Fig. S4), we generated 512 independent complete Emerson cohorts from the fixed Emerson-excluded model. Each donor’s released count total set its number of model draws: measured templates for 707 repertoires and one count per distinct amino-acid string for the other 61. Each simulation yielded *U*_*r*_*/D*_*r*_, where *U r* is the number of distinct strings private to donor *r* within that complete cohort and *D r* is its distinct-string count; both counts were random. We summarised each donor’s fraction and the across-donor interquartile range with simulation medians and 95% predictive intervals (Supplementary Methods S4). The donor intervals are pointwise, not simultaneous, and all intervals condition on the fixed model and draw counts, excluding model-fitting uncertainty. Private status here is defined within the held-out Emerson cohort; the descriptive comparison in Fig. 4b uses the full corpus.

The predicted plane of *P*_seq_ and publicness (Fig. 4d) gives, in columns of 0.1 decade of *P*seq, the expected number of sequences with each value of *m*, from the Poisson-binomial calculation at every bin of the spectrum. For this density, the corpus depths are approximated by 1,024 quantile groups, each represented by its geometric mean depth and its repertoire count. In the recorded comparisons with the ungrouped calculation, the measured total-variation discrepancy was 2.1 × 10^−3^. The analytic expected-publicness curve uses the ungrouped depths. It is also drawn on the observed plane (Fig. 4c) and gives the expected publicness of a sequence that is seen, E[*m* | *m* ≥ 1] =∑_*r*_ *θ*_*r*_ (*p*)*/*[1−∏ _*r*_(1 −*θ*_*r*_ (*p*))]. This conditional mean is a reference curve, not a prediction envelope or a significance threshold. For Supplementary Fig. S6a, length-16 sequences were divided into three bands of *P*_seq_, with cuts at 3.55×10^−10^, where the expected number of repertoires of the corpus carrying a sequence, ∑_*r*_ *θ*_*r*_ (*p*), equals two, and at 10^−6^. Each band’s probability-weighted positional composition is computed by banded dynamic programming on a grid of up to 8,192 bins and drawn as a sequence logo, as in Fig. 2h. These conditional logos describe sequences drawn from the model within a probability band; they do not assign equal weight to every possible string in that band. Band-restricted values inherit grid discretisation, whereas the unconditional logo is exact.

For the external publicness test (Fig. 4h; Supplementary Fig. S5a), we fitted one BEAM model per Rubelt population—naive and memory CD4 and CD8—to all 10 donors [18]. Clone-record weights, order four, equal directional weights, maximum length 27 and training-only prior estimation followed the family-held-out procedure above. The external source was AIRR Data Commons [20] dataset `IR-T1D-000001` at https://t1d-1.ireceptor.org, whose study record names Culina et al. [21] as its source, with 188 library records declaring 30 donors. For each donor and fraction, we selected the earliest eligible visit, breaking ties by source identifier, requiring at least 5,000 distinct productive standard-amino-acid CDR3 *β* strings of lengths 1–27. All recorded disease groups were retained. This yielded 29 CD4 CD45RA-positive, 30 CD4 central-memory, 29 CD8 CD45RA-positive and 20 CD8 central-memory catalogues. CD8 CD45RA-positive fractions include naive and terminal effector-memory cells; external central-memory fractions are narrower than the Rubelt memory groups. Importer-assigned record counts do not represent molecular abundances and were not used.

From each eligible library, we sampled 5,000 distinct amino-acid strings uniformly without replacement, using source-identifier-specific seeds derived by SHA-256 from seed 20261012. The first 2,500 entries defined nested smaller catalogues. Within each fraction, a string was public if present in at least two donors’ selected catalogues and private if present in exactly one. The primary evaluation excluded strings observed in any of the four training populations. Each string was scored under its corresponding population model and its independent-position control *Q* (Eq. (5)). The area under the receiver operating characteristic curve (ROC AUC) is the fraction of public–private pairs ordered correctly, giving ties half credit. Within-length AUC restricts these comparisons to exactly equal lengths and pools wins and pair counts across informative lengths. Donor AUCs were averaged equally within each fraction. Public/private labels and ranking statistics were recomputed for the nested smaller catalogues. Average precision and full-catalogue comparisons with length-only, training-sequence-frequency and training-donor-presence scores were also retained. Fractions within a donor and publicness labels across donors are dependent; no inferential intervals or tests were added. This test evaluates publicness ranking among observed catalogues in the four population-specific fits.

#### 3.9 Repertoire distance

The local-block Hellinger distance between two repertoires *A* and *B* compares their BEAM models through their length distributions and local sequence patterns. For each length *L*, let *b*= min (*k*+ 1, *L*), the span of the model’s longest factor, and let *P*^(*L,v*)^ be the exact marginal distribution, under *P* (· |*L*), of the block of *b* consecutive residues 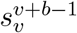 that starts at position *v*, for *v*= 1, …, *V*_*L*_ with *V*_*L*_ =*L*−*b*+ 1. With the squared Hellinger distance between distributions *µ* and 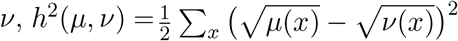,

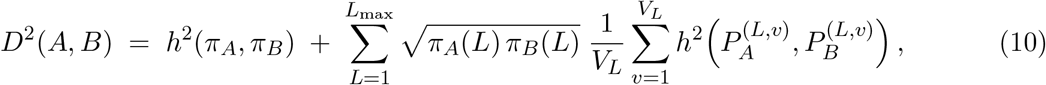

where *π*_*A*_ and *π*_*B*_ are the two models’ length distributions. *D* is the Hellinger distance between the joint distributions of a length, a block position drawn uniformly, and the residues of that block, so it is symmetric, satisfies the triangle inequality and lies between 0 and 1 (Supplementary Methods S6). All terms of Eq. (10) are non-negative and sum to *D* ^2^. The first term is the contribution of length frequencies; each term of the sum is the contribution of sequence patterns at one length and block position, which gives the decomposition of Fig. 5e. Block marginals are computed exactly by forward and backward sweeps, summing over all sequences allowed by each model.

#### Benchmark procedures

The 69 mouse repertoires (Fig. 5a) were clustered by Ward linkage [27]. The tree was cut into three clusters, whose labels were matched to the sorted populations by the assignment with greatest agreement. For the human naive–memory comparison (Fig. 5b), the original RDI clone tables were not recovered, so we used the reannotated cohort. We retained productive TRB records with standard-amino-acid junctions, complete unambiguous nucleotide codons and agreement between the source amino-acid annotation and its standard-code translation. We collapsed exact V-call/J-call/nucleotide-junction keys and excluded junctions longer than 27 residues. Distinct keys with identical amino-acid sequences remained separate unit-weight observations. Sample names and structured subset labels were supported by the original ENA library metadata and took precedence over conflicting free-text descriptions in 20 samples. Distances between naive and memory models were computed separately in CD4 and CD8 for each donor.

For the sampling comparison (Fig. 5c), each source repertoire was randomly split without replacement into disjoint subsamples of 100, 250, 500, 1,000 or 2,500 records, with five repeats per donor and size. A donor was included at a size only if both its naive and memory sources held at least twice that many records. Each subsample was fitted independently. The same-source distance averaged the naive–naive and memory–memory distances; the biological distance averaged the two orientations of naive–memory pairs at the same depth. Repeats were first averaged within donor, and the plotted lines and bands are donor medians and interquartile ranges. The primary size, 250 records, was specified before scoring. These splits estimate an operational sampling floor for finite observed pools, rather than an independent-assay variance.

Unequal-depth controls used disjoint subsamples of 100 and 400 clone records with the same 5 repeat seeds, source permutations and fitting procedure. Both sources were required to hold at least 500 records, retaining all 10 donors in both lineages. Same-source comparisons and both naive– memory size orientations used this depth pair. Supplementary Fig. S8a, b displays donor means over repeats, means of the two donors in each twin family, and individual repeat contrasts. These paired contrasts are descriptive.

For the longitudinal benchmark (Supplementary Fig. S10), day 0 denotes the donors’ return from travel; symptoms began on day 3. Earlier samples were collected one or two years before infection [24]. We first removed source records with empty or nonstandard amino-acid junctions, then selected up to 10,000 clone records per repertoire ranked by decreasing published `cloneCount`, preserving source order for ties. Separate nucleotide-defined records with the same amino-acid sequence remained separate unit-weight observations. Length filtering to 1–27 residues followed selection, excluding three records without replenishment. We computed nearest neighbours excluding the repertoire itself and clustered the full distance-matrix row profiles using Euclidean distance and complete linkage. Peripheral blood mononuclear cell (PBMC) distance summaries were restricted to PBMC–PBMC pairs. Mean PBMC distances were 0.128 between replicates, 0.154 across times within one donor and 0.264 between donors at the same time. These summaries describe the two sampled donors rather than estimating population-level temporal variability. Other methods applied to the sorted-population and naive–memory benchmarks (Supplementary Fig. S7) are described in Supplementary Methods S7.

For the larger human CD4/CD8 benchmark (Fig. 5d), we fitted order-four models to equal-weight distinct sequences and evaluated Eq. (10) over all lengths and block states. At each depth, every method received the same sampled sequences. Within-study reference sets excluded both fractions of the query donor; reciprocal transfer used only references from the other study. We evaluated mean within- and between-cell-type distances and nearest-neighbour accuracy, averaging the two fractions and three subsampling repetitions within donor. The depth comparison retained the same donors throughout. Intervals used 10,000 donor bootstrap draws conditional on the observed reference cohort (Supplementary Fig. S11; Supplementary Methods S7).

#### Controlled simulations

Five independent immuneSIM 0.8.7 repertoires were generated for the baseline and for each of four perturbations (Supplementary Fig. S9a): V/D/J usage noise (`vdj_noise`= 0.5), no N1 insertions, no 5^*′*^ D-gene trimming, and implantation of the three-residue motifs YAY and YVY into a combined 5% of sequences at random positions. All other generator parameters were defaults. Each replicate began with 12,000 generated sequences; after retaining standard-alphabet sequences of lengths 6–27 whose original amino-acid junction agreed with its nucleotide translation,10,000 records were sampled without replacement. The consistency check preceded motif modification, and BEAM used the modified amino-acid strings without requiring their modified nucleotide strings to agree. Each generated record received unit weight. The displayed value for a repertoire is its mean distance to the baseline repertoires, omitting self-comparisons. Synonymous codon replacements in baseline records supplied an input invariance control: identical amino-acid inputs give identical BEAM models.

#### Distance attribution

Fig. 5e decomposes the DN–CD4 squared distance for the mouse with the lowest numerical ID, chosen for illustration. Length and window contributions were computed for all 23 paired mouse repertoires from the terms of Eq. (10). Fractions for the illustrated pair use that pair’s *D*^2^ as the denominator; cohort and split shares divide each mean contribution by the mean *D*^2^ over the same paired mice. For pattern selection (Fig. 5f), a fixed random split reserved 11 mice and used 12 for discovery, keeping all populations of a mouse in the same group. Eligible lengths had at least five records in both DN and CD4 in at least eight discovery mice and were at least five residues long. Every five-residue pattern at each eligible length and window position was ranked by its mean contribution to the discovery DN–CD4 squared distance. Five windows were selected in descending order, requiring distinct strings and distinct length–position coordinates; this rule did not exclude overlap. The selected coordinates were locked before their observed within-length frequencies were compared between DN and CD4 in the reserved mice. Thus, mice were reserved for pattern selection, rather than for validation of the entire analysis. For each window and repertoire, observed frequency is the number of clone records matching that word at its specified position divided by all records of the selected length. Fig. 5f shows the paired DN–CD4 differences and their unweighted mean across reserved mice. Window contributions in both splits, depth and V-call controls, and the later complete-motif characterisation are reported in Supplementary Fig. S12 and Supplementary Methods S6. The complete-motif analysis followed inspection of the window results in these mice and is descriptive.

#### Statistical inference

The CD8-minus-CD4 naive–memory distance contrast (Fig. 5b) was averaged within each twin family and tested by exact two-sided sign-flip over the five families; the smallest attainable two-sided *P* is 0.0625. For the sampling comparison (Fig. 5c), the mean biological-minus-same-source gap at the primary size was calculated over family means, with delete-one-family jackknife *t* intervals at 97.5% nominal coverage in each lineage (95%simultaneous coverage for the two lineages by Bonferroni). Repeated splits were not treated as independent donors. Each simulation perturbation (Supplementary Fig. S9a) was compared with baseline by the mean between-condition distance minus half the sum of the two mean within-condition distances, excluding diagonal pairs. One-sided permutation *P* values enumerated all 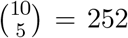 assignments of the ten repertoires to the two conditions; repertoires, rather than pairs, were the independent units. Each of the five originally selected windows (Fig. 5f; Supplementary Fig. S12a) was tested by exact one-sided sign-flip of the paired observed within-length frequency differences over reserved mice, in the direction established in discovery. Sign-flip inference assumes independent blocks and sign exchangeability under the null. Holm’s procedure [38] corrected the four simulation tests and the five window tests separately; the correction does not require the overlapping window tests to be independent. All five original windows were DN-enriched in every reserved mouse (Holm-adjusted *P* ≤ 0.00244). No additional confirmatory test was assigned to the post hoc complete motif. Clustering and nearest-neighbour results are descriptive.

### 3.10 Software

BEAM is implemented in C with OpenMP and has a Python interface. Models use the common specification and estimation procedure described above. Population and per-repertoire fitting times, peak memory, saved model sizes and distance-matrix scaling are reported in Supplementary Table S2. On one workstation (Intel Core i7-10700F, 16 threads), with the population reference model loaded, scoring 10^6^ sequences sampled from it took 1.0s on one thread and 0.16s on 16; the number of possible sequences took 1.16 s, the model’s entropy 2.75 s and the positional composition of one length 0.26s. The heaviest whole-space computations, the probability spectrum and the contour lines of the occupancy calculations, took minutes; contour values were cached and reused within each model. The BEAM analysis pipeline used Python 3.12 with NumPy, SciPy and pyarrow; the larger human CD4/CD8 model fits and exact distance calculations used Python 3.11. The SoNIA comparator used Python 3.11 with soNNia 0.4.0. Figures used R 4.3.3 with ggplot2 and patchwork.

## Data availability

The empirical TCR *β* repertoire data analysed in this study were obtained from the public sources cited in the Methods and Supplementary Information. Sources contributing to the population corpus are listed in Supplementary Table S1; corpus assembly, filtering and abundance weighting are described in Methods 3.1 and Supplementary Methods S1. The fitted BEAM models used in this study will be made publicly available at https://github.com/MuteJester/BEAM upon publication, allowing their use without downloading the full training corpus or refitting the models.

## Code availability

The BEAM source code, including its C implementation and Python interface, will be made publicly available at https://github.com/MuteJester/BEAM upon publication. A Python package will also be released through the Python Package Index (PyPI) upon publication, with installation instructions provided in the repository.

## Funding

## Supplementary Information

### Supplementary Methods S1

#### Corpus assembly

The corpus was assembled from six public sources (Supplementary Table S1; Methods 3.1) and reduced by a repertoire-level and a sequence-level cascade of filters.

**Supplementary Table S1:** Composition of the corpus by public source. Counts are of retained repertoires, after all filters.

| Public source | Studies | Repertoires | Share |
| --- | --- | --- | --- |
| immuneACCESS (Adaptive immunoSEQ) [28] | 270 | 28,384 | 73.7% |
| TCRdb 2.0 [29] | 135 | 6,334 | 16.4% |
| AIRR Data Commons [20] | 31 | 2,744 | 7.1% |
| mal-ID [30] | 1 | 550 | 1.4% |
| VDJbase [31] | 9 | 447 | 1.2% |
| Britanova et al. [32] | 1 | 79 | 0.2% |
| **Total** | **447** | **38,538** |  |

#### Repertoire filters

Repertoire filters.
| Rule | Removed | Remaining |
| --- | --- | --- |
| Acquired from source repositories | — | 55,423 |
| Emptied by the sequence filters | 786 | 54,637 |
| Fewer than 100 sequence records retained | 3,202 | 51,435 |
| Exact-duplicate content | 94 | 51,341 |
| Near-identical duplicates | 11,606 | 39,735 |
| Non-human | 1,197 | 38,538 |

The 786 emptied repertoires were immuneACCESS export summaries rather than clone tables, holding per-frame totals with an empty CDR3 field. The minimum of 100 applies to sequence records; after the records of each amino-acid sequence are merged, 66 repertoires hold fewer than 100 distinct sequences (at least 80).

##### Duplicate removal

Identical samples are often deposited under different accessions, so metadata matching does not find them. We compared all 1,317,923,470 pairs of the 51,341 repertoires exactly, on the Jaccard index of their CDR3 *β* sets, *J*(*a*, ∩ *b*) = |*a ∩ b*| */*( |*a*| + |*b*| − |*a* ∩*b*| ). Its distribution is bimodal, with a background mode below 0.05, a minimum near 0.45 and a separate population between 0.5 and 0.88. Metadata inspection showed that 87% of pairs above 0.5 are the same sample deposited in two repositories under the same sample name, mostly TCRdb 2.0 re-hosting immuneACCESS studies. One repertoire of each such pair was removed. This content screen addresses exact and high-overlap duplicates; samples of the same donor at substantially different depths need not have a high Jaccard index. A subsequent audit using exact sequence containment of at least 0.90 and overlap of at least 0.50 among the 200 most abundant clones identified 528 retained libraries as candidate duplicate deposits. These libraries remain in the frozen corpus analysed here. None of the identified or borderline candidate pairs involved the 768 Emerson held-out repertoires. Repertoires in this literature aggregate therefore should not be interpreted as independent individuals.

#### Sequence filters

Sequence filters.
| Rule | Distinct sequences removed |
| --- | --- |
| No amino-acid CDR3 (null or empty) | 37,464 |
| Length < 3 residues | 19,728 |
| $\geq 50\%$ stop codons | 64 |
| Non-standard characters | 50,262,483 |
| Length > 27 residues | 1,267 |

The alphabet test is by far the largest filter. The excluded characters are the stop codon *, the ambiguity codes `X` and `B`, the non-standard residue codes `O` and `U`, the gap character _, and a few stray punctuation and whitespace characters. This is a stop-free standard-alphabet criterion, not an assessment of rearrangement productivity: the reading-frame annotation is not used, so a rearrangement annotated as out of frame whose amino-acid CDR3 is spelled from standard residues is kept.

The upper bound of 27 residues was chosen from the measured length distribution rather than by convention. The main mode ends sharply at 27; beyond it the density falls by four orders of magnitude into a valley and then rises into two further modes near 58 and 80 residues. These are distinct populations at rough multiples of a CDR3 length, consistent with concatenated or misparsed junctions introduced before deposition, rather than the tail of one distribution. Applied after the alphabet test, which removes most of them already, the cutoff excludes a further 1,267 sequences.

##### Abundance units

The repositories report four kinds of count: immunoSEQ template counts (27,965 repertoires), sequencing reads (6,052), duplicate counts (3,434) and record counts (1,087). Template counts are platform-reported estimates of original input molecules, corrected using internal bias controls [33, 34]; they are not direct cell counts. Read counts retain PCR duplication: read-counted repertoires report 17 times more copies per distinct sequence, and summed together the reads repertoires (15.7% of the total) would carry 55.7% of the weight while contributing 10.4% of the sequence observations. Reads cannot be converted to templates, because no template counts were deposited for these repertoires. We therefore kept template counts where reported and assigned unit weight per distinct amino-acid sequence within each repertoire elsewhere, after merging records with the same amino-acid sequence. This minimum-count convention changes only the weights, not the sequences: the total observation weight is 5,629,439,400, compared with 19,974,430,818 summed counts in the original mixed units. The assigned unit weights indicate sequence presence and do not estimate missing template counts.

### S2 Exact normalisation

Notation is that of the Methods. Write 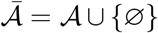 for the 20 residues and the boundary symbol, and for a string *x* of length *L* set *x*_*i*_ =*∅* for *i* <1 and *i* > *L*. The weight Φ _*L*_ (*x*) of Eq. (1) is a product of 2 *L* factors,

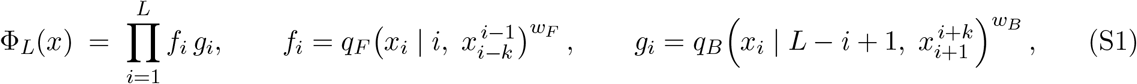

Where *fi* depends only on the window 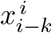 and *g*_*i*_ only on 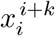 each spans at most *k*+1 consecutive symbols.

Consider the layered graph whose node at layer *j ∈* {0, …, *L*} is a state 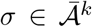, the last *k* symbols written, 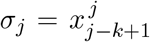. An edge from layer *j*−1to layer *j* appends the residue *x*_*j*_, so it knows the window 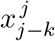. It carries the forward factor *f*_*j*_, whose window this is, and the backward factor *g*_*j*−*k*_ of the residue *k* positions back, whose window 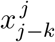 is now complete (with *g*_*i*_ ≡1 for *i* <1). The last *k* backward factors, whose windows extend past the end of the string, depend only on the final state and are collected in a terminal weight 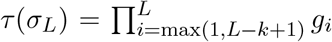 . Each string of length *L* is exactly one path from the initial state *∅*^*k*^ to layer *L*, and its weight Φ _*L*_ (*x*) is the product of its edge weights and its terminal weight. The forward and backward recursions

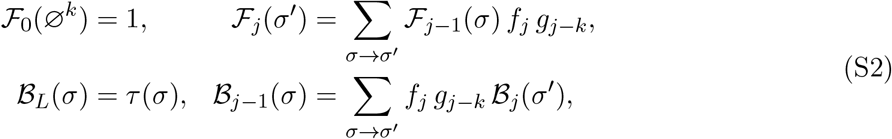

where the sums run over compatible edges into *σ*^*′*^ or out of *σ*, give

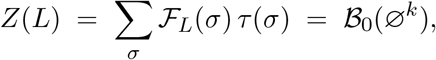

the sum of Φ*L* over all 20^*L*^ strings, in *O*(*L*·21^*k*^ ·20) operations. *ℱj*( *σ*)is the total weight of all prefixes that end in state *σ*at layer *j*, and ℬ*j*( *σ*) that of all completions from it. The marginal probability that positions *v*, …, *v*+*b*−1hold the residues *y* is therefore

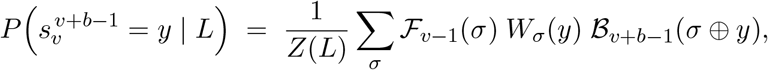

where *W*_*σ*_ (*y*) is the product of the *b* edge weights along the path that writes *y* from state *σ*, and *σ ⊕ y* is the state it ends in. With *b* = 1 this gives the positional marginals of Fig. 2h, and with *b* = min (*k*+ 1, *L*) the block marginals of the repertoire distance. The same expression applies to longer literal patterns, including the nine-residue event in Supplementary Fig. S12c; it sums over all compatible flanking sequences without assuming that overlapping windows are independent. These recursions use one-based sequence indices; the window starts displayed in the figures are zero-based.

These recursions evaluate the complete finite sums in floating-point arithmetic. They require no sequence-space sampling or pruning. With *Z* (*L*) computed once per length, Eq. (1) scores any string in *O*(*L*)table look-ups, and

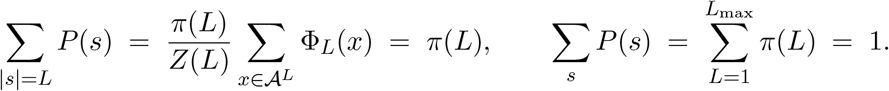

The normalisation holds for any tables and weights. The same exact individual-position marginals define the independent-position control *Q* in Eq. (5) (Figs. 2e and 4h; Supplementary Figs. S1a, b and S5a). Since each marginal sums to one, Σ_*s* =*L*_ *Q*(*s*) =*π*(*L*); *Q* therefore shares the fitted model’s length and individual-position distributions while removing between-position dependence.

Supplementary Fig. S3b reports the probability mass reconstructed from the counting bins.

### S3 Estimating the prior strengths

Consider one row (*t, u j*) of one expert at level *j*, with counts *c*(*t, u*_*j*_, *a*) and parent prediction 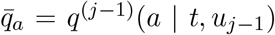 Give the row’s residue distribution *ψ*= (*ψ*_*a*_)_*a∈A*_ the Dirichlet prior with mean 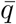 and concentration *α*_*j*_,*ψ ∼* Dirichlet 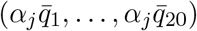, and treat the counts as multinomial given *ψ*. The estimated parent prediction 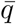 is held fixed in this conditional calculation. By conjugacy the posterior mean is

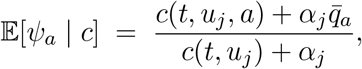

which is Eq. (3). Integrating*ψ*out gives the probability of the row’s counts,

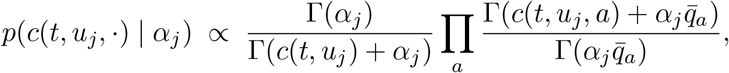

up to the multinomial coefficient, which does not depend on *α j*. Treating rows and observations as independent and summing the logarithm over the observed rows of the level, in both experts and at every position, gives Eq. (4). This rowwise score provides an approximate objective for strength selection; it is not the integrated likelihood of the normalised sequence model and does not integrate parent-table uncertainty. A residue with *c*(*t, u j, a*) = 0 contributes zero to the inner sum, and a row with no counts contributes zero in total, so the evidence runs over observed cells only. Counts weighted by abundance need not be integers; the expression is used as written, with Γ evaluated at real arguments.

The evidence of level *j* depends on the parents’ predictions, so the levels are estimated in order, *j* = 0,1, …, *k*, each with the strengths below it fixed. Two properties of the evidence shape the search. It can be flat: a row holding a single observation has the same evidence at every strength, so a corpus of very few sequences leaves *α*_*j*_ undetermined. The weak hyperprior log *α*_*j*_ *∼ N* (log 10,(log 100) ^2^), two decades either way at one standard deviation, regularises weakly informative estimates towards 10. Its relative influence decreases when the row evidence is informative. The evidence can also have several local maxima. The maximum of the evidence plus the Gaussian log density of the logarithmic strength is therefore located first on a grid of quarter-decade steps over [10 ^−3^,10 ^12^] and then refined by golden-section search within the best grid cell. The length strength *α*_*ℓ*_ is estimated in the same way, with the uniform distribution over 1, …, *L* max as the parent

### S4 Exact occupancy

#### Power sums

For any complex *z, P*(*s*) ^*z*^ =*π*(*L*) ^*z*^ *Z*(*L*)^−*z*^ Φ _*L*_ (*s*)^*z*^, and 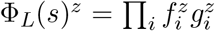 has the form of Eq. (S1) with every factor raised to the power *z*. The forward recursion of Eq. (S2) with powered factors therefore gives *Z z*(*L*) =∑ _|*x*|=*L*_ Φ*L*(*x*)^*z*^ exactly, and

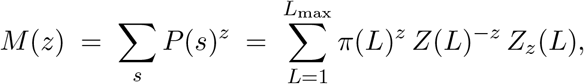

with *M* (0) the size of the model domain and *M* (1) = 1.

#### Occupancy quantities

For small depths the binomial expansion

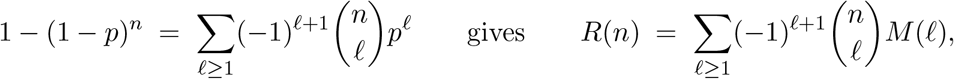

a finite sum for integer *n*, which converges quickly while *n* max*s P* (*s*) is small. For larger depths the Mellin–Barnes form is

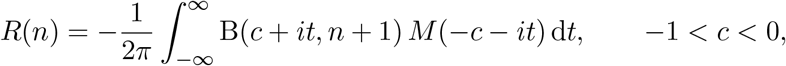

where B(*z, n*+ 1) = Γ(*z*)Γ(*n*+ 1)*/*Γ(*n*+ 1 +*z*) is the beta-function kernel. Other occupancy sums use the corresponding kernels or differences in the expected distinct-sequence count below. The integral is evaluated by the trapezoid rule on a truncated vertical line, using conjugate symmetry. The contour location, step size and truncation are chosen adaptively for each depth range; computed power sums are cached and reused.

The numerical implementation plans for relative tolerance 10^−10^ and records separate contributions from contour aliasing, tail truncation and floating-point rounding. For *R* (*n*), if *q*= −*c* and *h* is the integration step, the aliasing contribution is bounded using

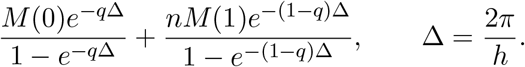

The tail contribution is evaluated from a numerical envelope of the kernel magnitude, and rounding accounts for the size of the kernel and dynamic-program phases. The reported relative bound is the total absolute bound divided by the computed magnitude minus that bound, when the latter is positive; otherwise no finite relative bound is certified. Values that do not meet the requested tolerance retain their achieved bounds. Thus, occupancy calculations avoid probability-spectrum discretisation, while their numerical accuracy remains explicitly recorded.

#### Sharing and private counts from expected distinct-sequence counts

The quantities of Figs. 3e and 4a, b, e, f reduce to the expected distinct-sequence count*R*. Shared sequences satisfy Eq. (8). For the private sequences of repertoire *r* in a cohort of total depth *T*,

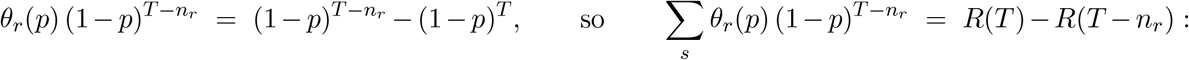

the expected private sequences of repertoire *r* are the sequences it adds to the rest of the cohort. The cohort’s expected private total is Σ_*r*_[*R*(*T*)−*R*(*T*−*n*_*r*_)], and for*N* repertoires of equal depth *n* each repertoire’s expected private sequences number *R* (*Nn*)−*R*((*N*−1)*n*).

#### Homogeneous-reference scenarios

Fig. 4e, f evaluates equal-depth cohorts under the same shared-distribution assumption. For *N* repertoires of depth *n*, a sequence is seen with probability 1 − (1− *p*)^*Nn*^ and is private with probability *N θ* (*p*) (1− *p*) ^(*N*−1)*n*^, where *θ* (*p*) = 1 − (1− *p*) ^*n*^. The predicted pooled private fraction is *N* [*R* (*Nn*) −*R*((*N* −1)*n*)]*/R*(*Nn*); the corresponding fraction within one repertoire is[*R*(*Nn*) −*R*((*N* −1)*n*)]*/R*(*n*). Across the plotted scenarios, private sequences remain a majority of the pooled catalogue, while their fraction within each repertoire decreases with cohort size. Numerous rare sequences occur in only one repertoire, whereas frequently sampled sequences recur across repertoires. At the observed depths, the ratio of mean private to mean distinct counts is lower than predicted in both the corpus and Emerson cohort.

Fig. 4g shows the publicness spectrum, expressed as distinct sequences per unit publicness in logarithmic bins: observed in the corpus; predicted at its recorded depths; predicted for the same number of repertoires at 10^11^ draws each; and predicted for a hypothetical cohort of 10^10^ repertoires at 10^11^ draws each. The corpus-depth prediction uses the Poisson-binomial calculation described above. For equal depths, *m*(*s*) is binomial with parameters *N* and *θ*(*p*). The corpus-sized deep scenario uses its exact binomial probabilities on the discretised sequence-probability spectrum; the largest scenario uses Poisson or normal approximations according to regime. Slopes were fitted by least squares on log–log axes between publicness 10 and 3,162; the approximate power-law decline over this range describes the plotted spectra and does not establish a universal scaling law. All scenarios condition on the fitted observational distribution, fixed draw counts and independent sampling. They illustrate the calculations across scales; extrapolation to heterogeneous human populations would require additional biological validation.

#### Predictive sampling intervals

Supplementary Fig. S4 evaluates the random private fraction *F*_*r*_ =*U*_*r*_*/D*_*r*_ in the 768-donor Emerson cohort [1], using the model built without those donors. We fixed each repertoire’s released count total as its draw count *n*_*r*_, retaining measured-template counts for 707 repertoires and sequence-presence counts for 61. These conventions define model draw counts and need not represent equivalent numbers of cells. Observed private and distinct counts were independently recounted by exact sequence matching across all donors in this cohort.

Each of 512 independent simulations generated the entire cohort. For each donor, we allocated *nr* draws multinomially over the model’s length distribution and sampled residues conditional on length using the backward recursion in Eq. (S2). At layer *j*, the transition from state *σ* to *σ*^*′*^ has probability *fj g*_*j*−*k*_*ℬ*_*j*_ ( *σ*^*′*^)*/ℬ*_*j*−1_ ( *σ*), so each choice accounts for all possible completions of the sequence. Exact sequence keys identified distinct strings within donors and shared strings across donors before computing *U*_*r*_*/D*_*r*_. Thus both counts varied between simulations, and generating complete cohorts retained dependence between donor fractions arising from the shared definition of private status. This distribution differs from the ratio of expected counts used in Fig. 4b. For each simulated cohort, we also computed the interquartile range of the donor fractions, using linearly interpolated sample quartiles; the observed cohort was summarised identically.

The replication count and independent random seeds were fixed before the production run. All production cohorts were retained; earlier pilot and diagnostic cohorts were excluded. For each donor fraction and for the cohort interquartile range, the 95% predictive interval uses the 12th and 501st ordered simulation values. For continuous exchangeable outcomes, these ranks give 489*/*513 marginal predictive coverage; ties are conservative. Donor intervals are pointwise. The number of observed donors outside their respective intervals is descriptive; donors were not treated as independent binomial observations.

Monte Carlo precision was assessed separately using exact binomial order-statistic 95%confidence brackets for the true 2.5th and 97.5th percentiles of the model’s sampling distribution (one-based simulation ranks 6–21 and 492–507, respectively). For the cohort interquartile range, these brackets were 0.988–0.999 and 1.073–1.084 percentage points. They describe uncertainty in estimated predictive endpoints, separately from sampling variation. Independent count recomputation and agreement of simulated mean counts with the analytical occupancy expectations checked the implementation. No model refitting was performed, and the intervals exclude uncertainty in the fitted model. Departures from the predicted distribution do not identify a biological or technical cause.

### S5 Expected entropy of a sample

Supplementary Fig. S3b compares model entropy *H* computed directly with its counting-spectrum estimate. The derivation below concerns the expected entropy of a finite sample.

Let *X*_*s*_ be the number of times sequence *s* is drawn in a sample of depth *n*. For a fixed-depth sample, the counts are multinomial and ∑_*s*_ *X*_*s*_ =*n*. The plug-in entropy of the sample is

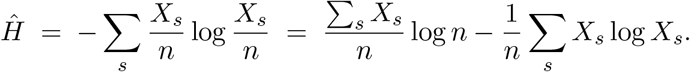

To approximate its expectation, we replace the counts by independent Poisson(*λ s*)variables with *λs* =*n P*(*s*). Their total is random, but E[∑_*s*_ *X*_*s*_] =*n*, so the first term still has expectation log *n*. All subsequent expectations in this derivation use this Poissonisation; the denominator remains the specified depth *n*. For *X∼* Poisson(*λ*)and any function *f*, E[*Xf*(*X*)] =*λ*E[*f*(*X*+ 1)], so E[*X*_*s*_ log*X*_*s*_] =*λ*_*s*_ E[log(1 +*X*_*s*_)]and

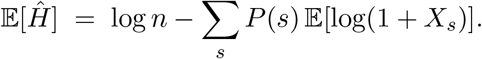

Since*H*=− ∑_*s*_ *P*(*s*) log *P*(*s*) = log *n*− ∑_*s*_ *P*(*s*) log*λ*_*s*_,

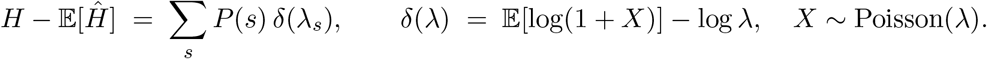

By Jensen’s inequality E[log(1 +*X*)] ≤ log(1 +*λ*), and*δ*(*λ*) is large for *λ≪* 1 and tends to zero as *λ* → ∞, so the shortfall comes from sequences expected to be drawn less than about once. Fig. 3d evaluates it by decade of *P*_seq_ on the spectrum. Its total is evaluated by a power-sum contour calculation with recorded numerical bounds, without spectrum discretisation (Supplementary Methods S4).

### S6 The repertoire distance

For a model with length distribution *π* and block marginals *P*^(*L,v*)^, define the joint distribution of a length *L∼π*, a block start *v* uniform on1, …, *V*_*L*_, and the block’s residues *y ∼P*^(*L,v*)^:

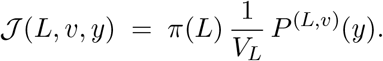

The Bhattacharyya coefficient 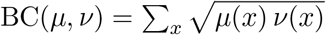 satisfies *h*^2^(*µ, ν*) = 1−BC(*µ, ν*). Because the distribution of *v* given *L* is the same for both models, the coefficient of *J*_*A*_ and *J*_*B/*_ factorises,

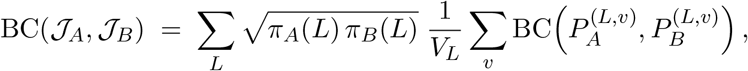

and therefore

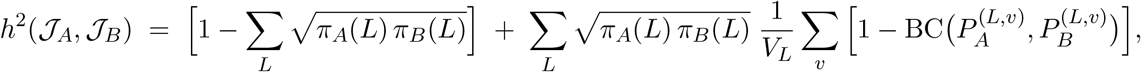

whose first term is *h*^2^(*π*_*A*_, *π*_*B*_). This is Eq. (10):*D*=*h*( *J*_*A/*_, *J*_*B*_). As a Hellinger distance between distributions, *D* is symmetric, satisfies the triangle inequality, lies in [0,1], and is zero exactly when *J*_*A*_ = *J*_*B*_, that is, when the two models agree on lengths and on every block marginal. For arbitrary sequence distributions, comparing these block marginals defines a pseudometric: differences that leave all of the chosen marginals unchanged are invisible to *D*.

The terms of Eq. (10) are non-negative and sum to *D*^2^. For *D* >0, dividing each contribution by *D*^2^ gives its share of the squared distance. The length-frequency share is *h*^2^(*π*_*A*_, *π*_*B*_)*/D*^2^; the share for sequence patterns at length *L* and block position *v* is

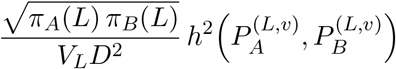

(Fig. 5e). The share of one pattern *y* is the corresponding summand of 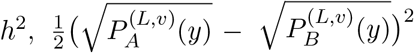, weighted in the same way (Supplementary Fig. S12e). The block marginals are computed exactly by the forward and backward recursions of §S2, so every term includes the sequences that neither repertoire contains.

Compatible overlapping windows can be grouped without changing the distance. Writing the contribution of a window pattern explicitly as

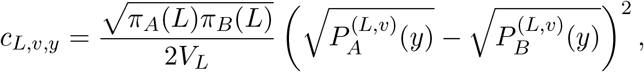

a group *G* has contribution 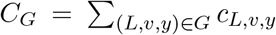. Each original coordinate is counted once; unreported coordinates remain in the remainder of *D*^2^. Overlap among the underlying sequence events does not invalidate this accounting, because these are additive terms of the defined metric. A group need not correspond to a disjoint set of receptors or an independent biological feature.

The complete sequence motif implied by compatible windows is a separate event: a sequence must satisfy all of the windows simultaneously at the specified length and positions. Its probability is evaluated directly with the constrained recursion of §S2; it is not the sum or product of the overlapping-window probabilities. Likewise, *C*_*G*_ is not the contribution of this complete event to a redefined distance. The five discovery-selected windows of Fig. 5f form one group aligned to `TISNERLFF` at zero-based positions 6–14 of length-15 CDR3s (Supplementary Fig. S12f). A receptor matching only some group members need not contain the full motif. Their summed original contributions account for 2.03% of mean reserved-mouse *D*^2^, calculated as mean grouped contribution divided by mean *D*^2^ over the same mice (Supplementary Fig. S12e). This quantifies one component of the distributed repertoire difference, rather than the contribution of the literal nine-residue event.

Distance matrices reuse each repertoire’s exact block marginals. For *M* repertoire models and *F* probability entries across the compared block distributions, all pairwise comparisons require *O*(*M*^2^*F*) operations after marginal extraction. The implementation processes one length at a time and forms pairwise Gram matrices one window at a time; the final dense distance matrix requires *O*(*M*^2^) storage. The quadratic term describes pairwise arithmetic; wall times also include data movement and other overhead. Supplementary Table S2 separates marginal preparation from matrix assembly.

#### Window controls and motif characterisation

The original windows were checked after 200 within-length subsamples per reserved mouse, drawing each population down to the smaller length-specific count without replacement, and after 3 full-repertoire depth-matched BEAM refits per reserved mouse. A descriptive within-V comparison used only V-call strata present in both populations at the selected length, weighted by pooled stratum counts within mouse; strata without common support were excluded. These controls assess depth and gene-composition sensitivity and do not establish a causal source of the pattern differences (Supplementary Fig. S12a, b).

After inspecting the original window results in the reserved mice, we evaluated the complete nine-residue motif directly by constrained dynamic programming, conditional on length 15. Its observed frequency counts records matching every residue of the complete event, divided by all records of that length in the same repertoire. Each mouse remained the paired unit, with equal mouse weights in summary contrasts. Observed frequencies and model probabilities describe the same fitted repertoires. This complete-motif analysis is a post hoc characterisation of already examined mice, not a separately reserved validation or an additional confirmatory test.

The complete motif was DN-enriched in all 11 mice in both observed within-length frequencies and fitted conditional probabilities, with mean DN–CD4 differences of 13.3 and 11.1 percentage points, respectively (Supplementary Fig. S12c). Its observed matches coincide with those of the original `TISNE`window, although their model probabilities differ. This follow-up therefore characterises the original evidence rather than supplying independent confirmation.

For the complete motif, we additionally counted each distinct amino-acid CDR3 string once within each repertoire and repeated the within-V standardisation. Identical observed membership allowed its equal-length-count control to use the original `TISNE` subsamples. We queried the complete event in each saved matched-depth model, rather than substituting its constituent-window probabilities. Enrichment persisted after distinct-CDR3 counting and in the matched-depth model means in all 11 mice, and within shared V-call strata in 10 (Supplementary Fig. S12c).

Carrier descriptions count distinct CDR3 strings per mouse and population; pooled V-call frequencies use the original matching input records. Pooled distinct-string and V-call totals are deduplicated across mice. Carriers span multiple CDR3 strings and V calls (Supplementary Fig. S12d). The sequence overlaps the reported Tremont-associated motif [22, 17]; this observation does not establish antigen specificity or causality. V calls do not define clonotype families, and no J-gene or antigen assignments were made.

### S7 Repertoire-distance benchmarks and comparators

The benchmarks of Figs. 5a and b were repeated with four other distance methods (Supplementary Fig. S7), each run from its authors’ code with the settings below; inputs and audits are recorded with the analysis. **TCRdist with optimal transport** [16, 17]: the code of Olson et al. supplied pairwise TCRdist costs using the mouse and human reference databases for the respective cohorts. Clone records had uniform weight. Costs were divided by 200 before applying `POT 0.9.7.post1 sinkhorn2`, with regularisation 0.01, at most 1,000 iterations and stopping threshold 10^−9^; the returned transport cost was multiplied by 200. This is the transport term under an entropically regularised plan, without a self-cost correction. Mouse diagonal entries were set to zero for clustering. Pairs exceeding the stopping threshold were additionally examined with up to 50,000 iterations and threshold 10^−12^; primary comparisons retain the original settings. For human gene calls without alleles, the reference allele *01 was used. Equivalent multi-gene calls were represented by their first candidate; 147 of 151,625 rows were omitted (91non-equivalent multi-gene calls and 56 additional CDR3s of five residues or fewer). **Repertoire Dissimilarity Index** [5]: the authors’`calcRDIon` V-gene calls, at a common depth of 68(mouse) or 610(human) clone records, with 100 repeats. **immuneREF** [35]: four of its six features, amino-acid composition, amino-acid architecture, aminoacid overlap and V-gene usage, with equal weights, since the tables lack the nucleotide, D, J and abundance fields the others need; the authors’ code does not count multi-gene V calls (3,477 human rows). **CompAIRR** [39]: exact amino-acid matching of unique sequences, with the Jaccard distance 1−|*A*∩ *B*| */* |*A ∪B*| . Mouse trees were built with the same Ward [27] display for every method. These configurations are reduced or exact-only forms of immuneREF and CompAIRR, and CompAIRR compares sets of unique sequences whereas the other methods weight clone records, so the comparison is descriptive.

#### Larger human CD4/CD8 benchmark

Fig. 5d shows the primary reciprocal-transfer comparison of BEAM with its same-fit positional projection. Supplementary Fig. S11 reports the full evaluation of cell-type separation, donor-held-out matching, input depth and reciprocal study transfer under a design fixed before scoring. GSE285951 [25] provides 62 RNA/UMI blood TCR libraries, of which 60 form complete CD4/CD8 pairs from 30 people; two people with CD4-only libraries were excluded. The paired set comprises 24 people with head and neck squamous cell carcinoma, 1 with ameloblastoma and 5 healthy donors. We used the deposited MiXCR TRB clone tables from SMART-Seq RNA sequencing with unique molecular identifiers. The immunoSEQ cohort comprises 44 CD4/CD8 libraries from 22 people with multiple sclerosis [26]. Person identities were recovered from the shared subject identifiers in original sample names and reconciled with the cell-subset metadata; six additional exports had no productive sequence records. No exact catalogue duplicates or cross-source pairs with at least 95% containment or Jaccard similarity were found.

We retained non-empty CDR3 *β* strings of length 1–27 containing only standard amino acids and collapsed identical amino-acid strings within each library. Emerson records were required to carry the source productive-frame flag; no additional translation-based filter was applied to the deposited GSE tables. Every distinct string received unit weight, irrespective of its read, UMI or template count. For each of 3 repetitions, a seeded permutation of the sorted catalogue supplied nested prefixes of 1,000, 2,500, 5,000 and, where available, 10,000 strings. PCG64 seeds were the first eight bytes, read little-endian, of the SHA-256 digest of `study70:source:source_sample_id:replicate`, with repetition indices 0–2. The primary 1,000-sequence evaluation used all paired donors. The main depth comparison fixed 26 GSE and 22 immunoSEQ donors at all three depths (Supplementary Fig. S11c, d). A separately reported 10,000-sequence extension fixed 18 and 22 donors across all four depths. Thus, depth effects do not reflect changing donor sets.

Models used order four, maximum length 27, equal forward/backward weights and the standard evidence-fitted smoothing. Eq. (10) was evaluated over every length, position and block state, without truncating low-probability terms. The same-fit positional projection retained the fitted models’ length and individual-position marginals, comparing the joint distribution *Q*_*p*os_ (*L, i, a*) = *π*(*L*)*P*(*s*_*i*_ =*a* | *L*)*/L*by Hellinger distance. Each position has equal weight within its length. This control compares exact marginals from the same fit, rather than refitting an independent model or using empirical residue frequencies. Empirical Hellinger controls compared observed length distributions, joint length–position–residue distributions and joint length–window–block distributions, using the same uniform position/window convention and blocks of min (5, *L*) residues. CompAIRR [39] supplied exact amino-acid Jaccard dissimilarities, independently checked against set intersections. The V/J control used Hellinger distance on concatenated V- and J-gene frequency vectors, each carrying total mass one-half. RDI [5] used the authors’ implementation on fractional V-gene profiles with `subsample=FALSE,constScale=TRUE` and `units=‘lfc`’, an explicit adaptation for ambiguous gene calls.

For the gene controls, each selected amino-acid string contributed one unit divided among its distinct gene options after allele removal and nomenclature harmonisation. For scored MiXCR calls, only maximum-scoring genes, including ties, were retained per source row; options were unioned over rows encoding the same amino-acid string. Unscored declared alternatives were retained. Unassigned calls remained an explicit category, including when assigned and unassigned rows encoded the same string. Mean unassigned V-gene mass at the primary depth was 0.0% in GSE and 20.6% in immunoSEQ. Gene-based methods therefore use additional information and differing annotation resolution; their performance is interpreted for these tasks, not as a universal ranking of repertoire distances.

Within-study queries excluded both CD4 and CD8 repertoires from their own donor. Transfer used reference and query repertoires from different studies, matched by depth and repetition, without target-label calibration. Mean distances to same- and opposite-cell-type references supplied Supplementary Fig. S11a. For nearest neighbours, numerical ties (absolute tolerance 10^−12^, relative tolerance 10^−10^) received the average correctness over tied references. A predefined secondary classifier selected the cell type with the smaller mean reference distance, with half credit for ties. The complete tables also report the probability that an opposite-type reference is farther than a same-type reference, with half credit for ties. This separation AUC summarises distance ordering, rather than a classifier ROC curve.

Each donor contributed equally after averaging both cell-type queries and all three repetitions. We used 10,000 fixed-seed donor bootstrap draws for 95% percentile intervals and paired method differences. These intervals describe query-donor variation conditional on the observed reference cohort; they do not resample reference cohorts. All-perfect donor scores produce a degenerate bootstrap interval, not a guarantee of perfect population performance. All predefined methods, both classifiers and depth sets are retained in the accompanying source-data tables. Sequence subsamples and repertoire pairs were not counted as independent people. Cross-study transfer combines changes in assay, processing and clinical cohort, so it does not isolate an assay effect. The labels assess CD4/CD8 matching, not disease diagnosis or naive–memory discrimination.

### S8 Sensitivity to model design and corpus weighting

Supplementary Fig. S2 evaluates prespecified changes to the model and its input weights using common observations for every configuration. The comparison characterises sensitivity rather than selecting hyperparameters on the evaluation cohort.

#### Corpus models

All 768 Emerson repertoires [1] were excluded before fitting or estimating prior strengths, leaving 37,770 training repertoires. The primary configuration used *k* = 4,(*w*_*F*_, *w*_*B*_) = (1*/*2,1*/*2) and the corpus weights: template counts where reported and one occurrence per distinct amino-acid string within a repertoire otherwise. The eight alternatives comprised order 2 or 3; directional weights (1,0) or (0,1); one occurrence per string in the entire corpus or per repertoire; and restriction to the same 27,258 template-counted repertoires, using either counts or within-repertoire presence. Global deduplication removes both abundance and cross-repertoire recurrence weighting. The template-only pair isolates weighting on identical sources; comparison with the primary model also changes training size and composition. Order and direction remained at their primary values in the weighting controls. All fits used standard-amino-acid release inputs up to length 27, without an additional in-frame filter, and prior estimation using only the fitting data (§S3). Order 5 was excluded in advance because the amino-acid implementation supports orders up to4.

#### Marginal fidelity

For Supplementary Fig. S2a, model length and residue marginals were computed exactly and compared with the original in-frame source-row histograms, including repeated amino-acid strings. Length histograms retain source lengths through 40; model probabilities are zero above 27. Overall composition counts standard residues, and positional composition pools lengths separately at each of the first 18 positions. Jensen–Shannon distances use natural logarithms, with the positional score averaging distances across positions as in Methods 3.6.

#### Sequence probabilities and recurrence

For each eligible donor, we sampled 5,000 distinct productive standard-amino-acid strings of lengths 1–27, uniformly without replacement from the lexicographically sorted catalogue. Productivity required `frame_type=In`. NumPy PCG64 was seeded using the first eight SHA-256 bytes, interpreted little-endian, of `BEAM-68-20261012:` followed by the repertoire identifier. One donor had insufficient eligible strings, leaving 767 fixed catalogues; it remained in the fidelity and accumulation analyses. The primary scoring subset excluded every string present anywhere in the full non-Emerson training union, including when evaluating template-only models. For each donor we averaged −log*P*(*s*) and −log*P*(*s*)*/* |*s*| over this subset, including the model’s length probability. Supplementary Fig. S2b shows the latter score minus the primary model’s score, paired within donor and then averaged equally across donors. Complete-catalogue scores were retained as a sensitivity analysis; the unseen subset represents a different evaluation distribution from the complete catalogue.

A string was public if present in at least two sampled donor catalogues and private if present in one. ROC AUC assigns half credit to tied probabilities. AUC and average precision required at least ten strings per class; 0 donors met this requirement in the unseen subset. Supplementary Fig. S2c therefore displays the prespecified secondary complete-catalogue AUC, which includes training-overlapping strings (76.9% of sequence–donor entries). No eligibility threshold or catalogue was changed. Average precision, within-length AUC weighted by the number of public–private pairs, and Spearman rank correlation with the primary model were also retained. Recurrence ranking does not establish absolute probability calibration.

#### Accumulation

Full held-out repertoires were pooled in 30 PCG64 donor permutations, using seed20260816and an identifier-sorted starting roster. The cumulative depth sums within-donor distinct-sequence counts; the final depth is 140,840,439, with 73,245,676 strings in the observed union. Expected distinct counts were evaluated at identical depths for every model at 1, 2, 4, 8, 10, 16, 32, 64, 100, 128, 256, 512 and 768 donors. Supplementary Fig. S2d shows mean predicted/observed ratios at 10,100 and 768 donors; all 13 checkpoints were retained. The occupancy calculation (§S4) requested relative tolerance 10^−8^, accepting results only with reported relative bounds at most 10^−6^. No model-specific depth rescaling was fitted.

#### Repertoire distance

The 40 processed Rubelt repertoires [18] described in Methods 3.9 were fitted separately under 6 configurations: bidirectional orders 2, 3 and 4, order-4 forward-only and backward-only models, and bidirectional order 4 after amino-acid deduplication within each repertoire. All original nucleotide-key records were retained without subsampling in the first five configurations. These fits used prior estimation from each repertoire and maximum length 27, independently of the corpus models. The comparison block width remained min(5, *L*)for every order, with all positions and lengths included, so changing order did not change the distance definition (§S6). Exact block marginals were independently checked against masked-probability queries and exhaustive enumeration on small models. For each donor we subtracted the CD4 naive–memory distance from the CD8 naive–memory distance; family means average the two twins (Supplementary Fig. S2e).

All configurations and eligible observations were retained. Red bars in Supplementary Fig. S2a– d span interquartile ranges across donors or pooling orders, not standard deviations or confidence intervals. Repeated configurations, overlapping pools and twins were not treated as independent biological replicates.

**Supplementary Table S2:** Computational cost of model fitting and exact repertoire distances. (a) A full population-model fit and representative single-repertoire fits at four input depths, measured in distinct CDR3 *β* amino-acid sequences. Per-repertoire entries are medians [minimum– maximum] across four fits at each depth; the population fit was measured once. (b) Exact block-marginal preparation for 32 repertoire models, each built from 5,000 distinct sequences, followed by distance-matrix assembly for nested sets of 8, 16 and 32 models using those marginals. Each assembly size was measured once.

| **a. Fitting input** | **Fit time (s)** | **Peak RSS (GiB)** | **Model (MiB)** |
| --- | --- | --- | --- |
| Population: 926,909,993 strings | 815.36 | 2.665 | 84.955 |
| Repertoire: 1,000 strings | 19.66 [19.46–20.39] | 0.138 [0.138–0.139] | 0.100 [0.097–0.102] |
| Repertoire: 2,500 strings | 19.75 [19.42–19.98] | 0.146 [0.145–0.146] | 0.213 [0.207–0.220] |
| Repertoire: 5,000 strings | 20.11 [19.52–21.04] | 0.154 [0.153–0.156] | 0.365 [0.350–0.380] |
| Repertoire: 10,000 strings | 20.20 [20.18–20.33] | 0.166 [0.165–0.169] | 0.606 [0.587–0.630] |

| **b. Distance computation** | **Models** | **Pairs** | **Time (s)** |
| --- | --- | --- | --- |
| Model loading and worker startup | 32 | — | 53.06 |
| Exact block marginals and scratch writing | 32 | — | 194.38 |
| Matrix assembly from cached marginals | 8 | 28 | 56.49 |
| Matrix assembly from cached marginals | 16 | 120 | 98.39 |
| Matrix assembly from cached marginals | 32 | 496 | 175.11 |

All models used order four, maximum length 27, equal directional weights and automatic prior-strength estimation. Fitting includes weighted counting, prior estimation and exact normalisation. The population run used the prepared, aggregated corpus Parquet table and includes reading and decompression; it excludes corpus assembly and input-hash verification. The input was read for verification before timing, without flushing the operating-system cache. Per-repertoire inputs were loaded before timing; these fits used paired CD4/CD8 repertoires from one prespecified donor per source in Supplementary Fig. S11. Saving and output verification are excluded from fit times. Model size is the size of the saved BEAM file. Peak RSS is the operating system’s maximum resident set size, including the interpreter and input buffers; population RSS includes saving as well.

Measurements used an Intel Xeon Gold 6252 server (24 physical cores, 48 hardware threads), Python 3.11 and the same frozen BEAM implementation as the larger human distance benchmark. The population fit used 16 BEAM threads and two Arrow threads; single-repertoire fits used one thread. Distance preparation used 16 one-thread workers, and matrix assembly used 16 BLAS threads. Preparation includes all lengths 1–27 and all block positions. Assembly includes reading/stacking cached roots, pairwise arithmetic and numerical checks; it excludes fitting and marginal preparation. The three assembly sizes were timed in ascending order, share one preparation run and are not independent end-to-end timings.

Across the combined distance run, the maximum sampled sum of resident memory over the parent and worker processes was 19.79 GiB (50-ms sampling; shared pages counted per process), and the maximum temporary block-root storage in RAM-backed scratch space was 17.55 GiB. Scratch pages can also be represented in RSS. This memory measurement applies to the shared run, not separately to each matrix size. One MiB is 2 ^20^ bytes and one GiB is 2 ^30^ bytes.

**Supplementary Figure S1:**
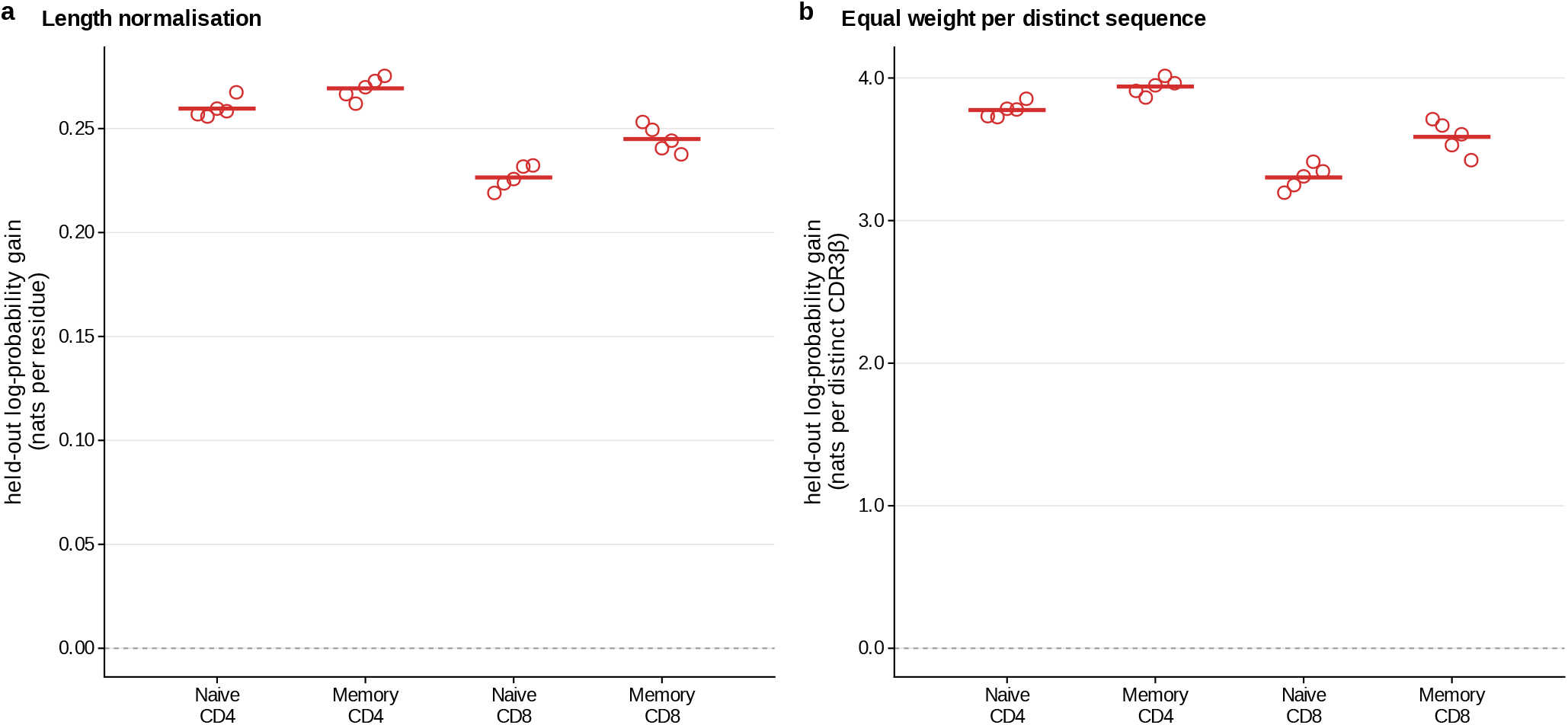
Length and sequence-weighting controls for held-out prediction. The same family-held-out fits and unseen-sequence comparison as Fig. 2e are evaluated with two scoring controls. Reannotated Rubelt repertoires [18] comprise 10 donors in 5 twin families and 4 sorted populations. A separate BEAM model for each population is fitted after excluding both twins from one family, using the other 8 donors; this gives 20 fits. The normalised independent-position projection Q preserves each fit’s length distribution and every individual-position marginal exactly. Test strings are absent from all populations of every training donor. (**a**) Mean natural-log probability gain, log[*P*_seq_(s)/Q(s)], divided by each sequence’s length before averaging. Each nucleotide-defined test record retains unit weight. (**b**) Mean log-probability gain with each distinct amino-acid string counted once within its test repertoire. Models and test membership remain the same in both controls. Each open point averages the two held-out donor means within one family and population; horizontal bars average families equally. The dashed line marks zero gain. Averaging families and populations equally gives 0.25 nats per residue in a and 3.65 nats per distinct sequence in b. Underlying donor–population means are positive in all 40 comparisons in a and all 40 comparisons in b. Molecular abundances are unavailable; nucleotide-defined records and distinct amino-acid strings represent different observation units. Training folds overlap and populations within a donor are dependent; no inferential intervals or tests are shown. These controls do not separate correlations associated with V/J genes from other residue relationships.

**Supplementary Figure S2:**
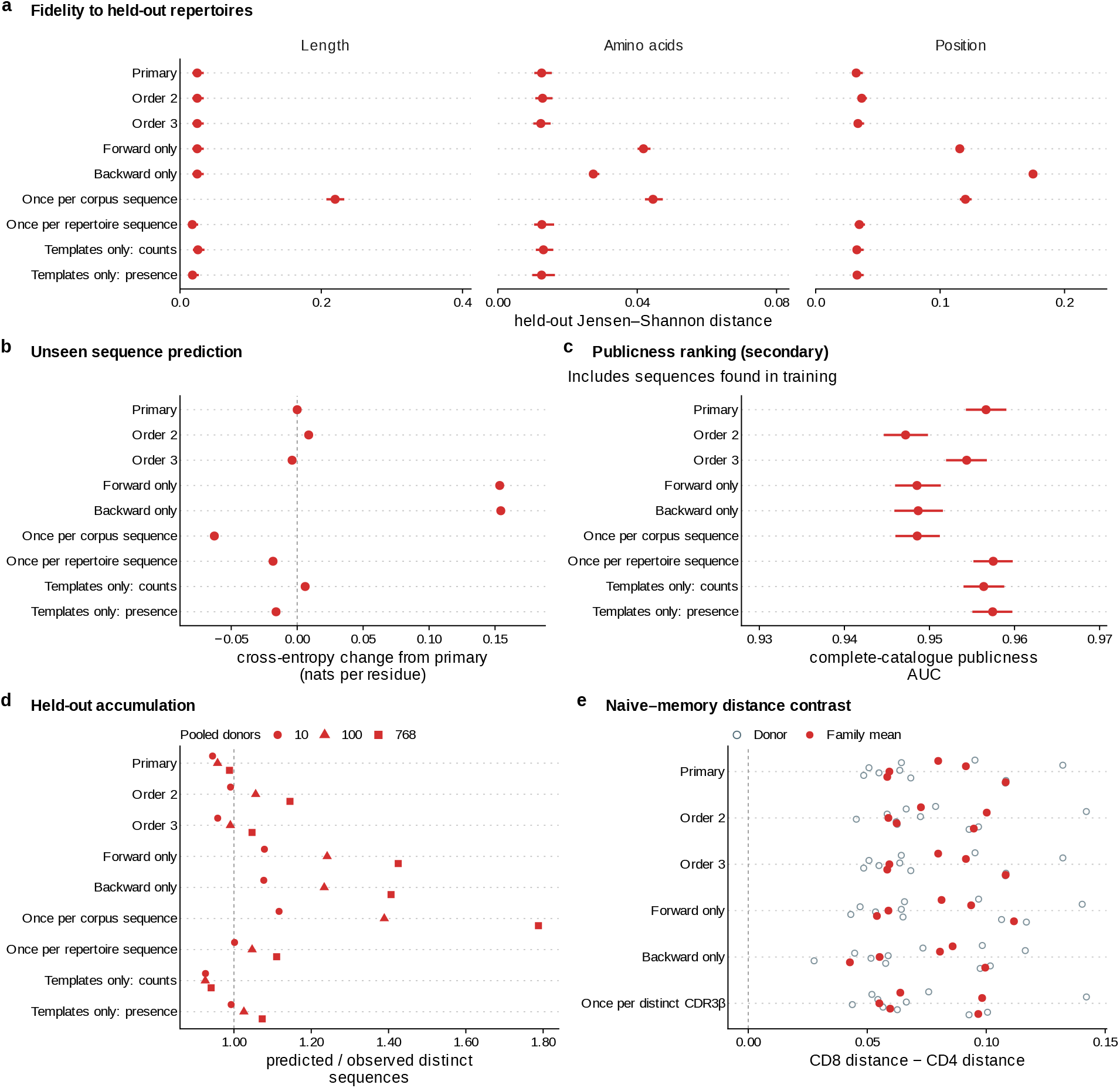
Sensitivity to model design and corpus weighting. 9 corpus models exclude Emerson donors [1]. Primary uses order 4, equal directional weights, template counts where available and unit weight per distinct sequence within each repertoire otherwise. Alternatives change order, direction or weighting. Once per corpus sequence removes abundance and recurrence weighting; once per repertoire sequence retains recurrence. Template-only rows restrict identical sources to counts or presence. (**a**) Held-out marginal fidelity across 768 donors: median length and amino-acid Jensen– Shannon distances and median donor mean distance across the first 18 positions. Lower is better; facets have separate scales. (**b**) Mean paired cross-entropy change per residue versus primary, for strings absent from the training union. Each of 767 donors supplies a uniform 5,000-string catalogue; only unseen strings are scored. Negative is better. Primary cross-entropy: 1.701 nats per residue (26.804 per sequence). (**c**) Mean donor ROC AUC ranking public above private strings in complete catalogues; public means detected in at least two donor catalogues. Chance is 0.5. This prespecified secondary analysis includes 76.9% training-overlapping sequence–donor entries. No donor met the primary unseen-ranking minimum of ten strings per class. (**d**) Predicted/observed distinct-sequence ratio after pooling 10, 100 or 768 donors in 30 orders. All models use identical full-repertoire sequence-presence depths; one is agreement. (**e**) CD8 minus CD4 naive–memory distance in 10 Rubelt donors [18] (open circles), with 5 twin-family means (red points). 6 configurations fit each of 40 repertoires using identical nucleotide-key inputs. All retain five-residue observation windows (complete sequences below length five); the final row counts distinct CDR3*β* once per repertoire. Positive means greater CD8 naive–memory distance. In a–d, red centres are medians (a) or means (b–d); red bars span the interquartile range across donors (a–c) or pooling orders (d), not standard deviations or confidence intervals. Donors/orders are reused across configurations; family means summarise the same donor observations.

**Supplementary Figure S3:**
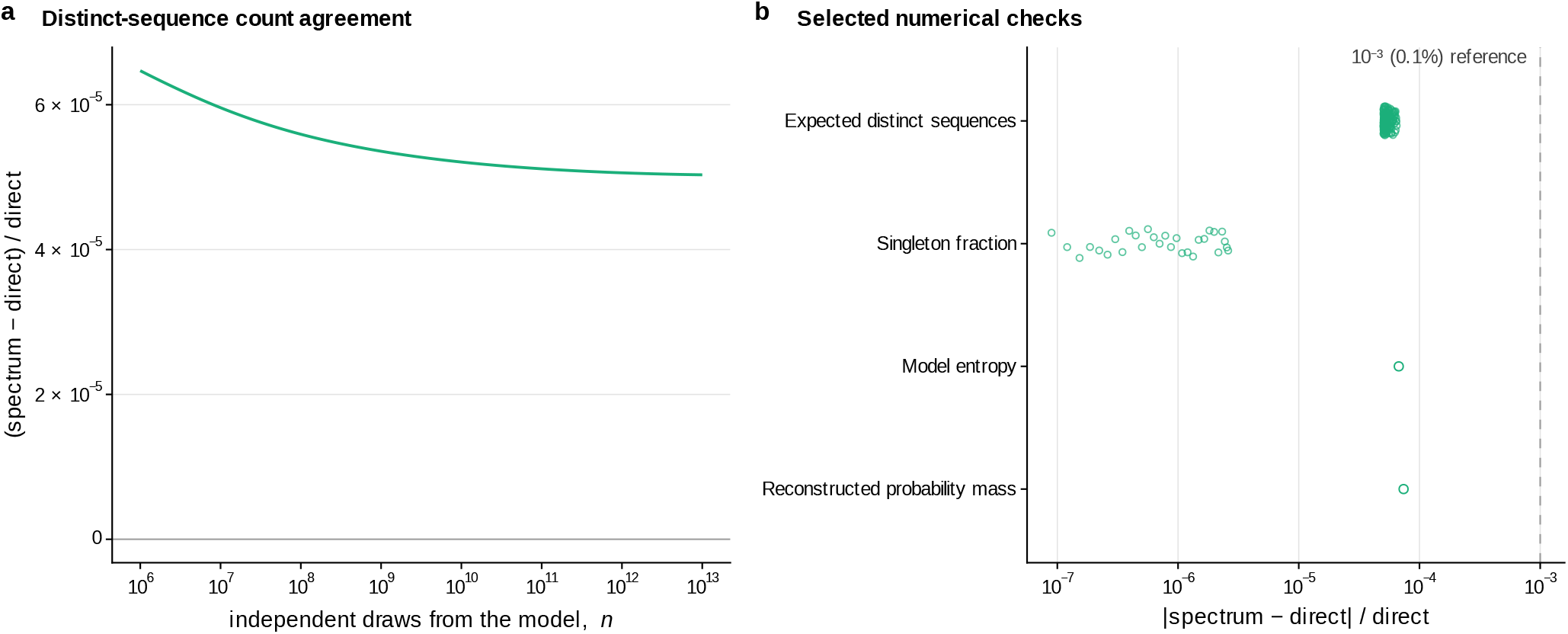
Numerical agreement of selected whole-space quantities. Selected sums over all sequences of one population reference model, computed directly (BEAM’s dynamic programs and spectrum-free occupancy calculations, with recorded numerical error estimates for occupancy) and on the 32,768-point counting spectrum used in Figures 2–4. (**a**) Relative difference, spectrum minus direct, in the expected number of distinct sequences R(n) at 141 depths from 10^6^ to 10^13^ independent draws; maximum absolute relative difference 6.5 *×* 10^−5^. Differences are divided by the direct value and shown as fractions in both panels. (**b**) Absolute relative differences in R(n) at the same 141 depths, the singleton fraction at 29 depths, model entropy, and probability mass reconstructed from the counting bins and their representative probabilities. BEAM’s sequence distribution is normalised to one. Points with several depths are spread vertically for legibility. Dashed line, 0.1% relative difference, shown as a reference. The largest absolute relative difference over all displayed comparisons is 7.4 *×* 10^−5^.

**Supplementary Figure S4:**
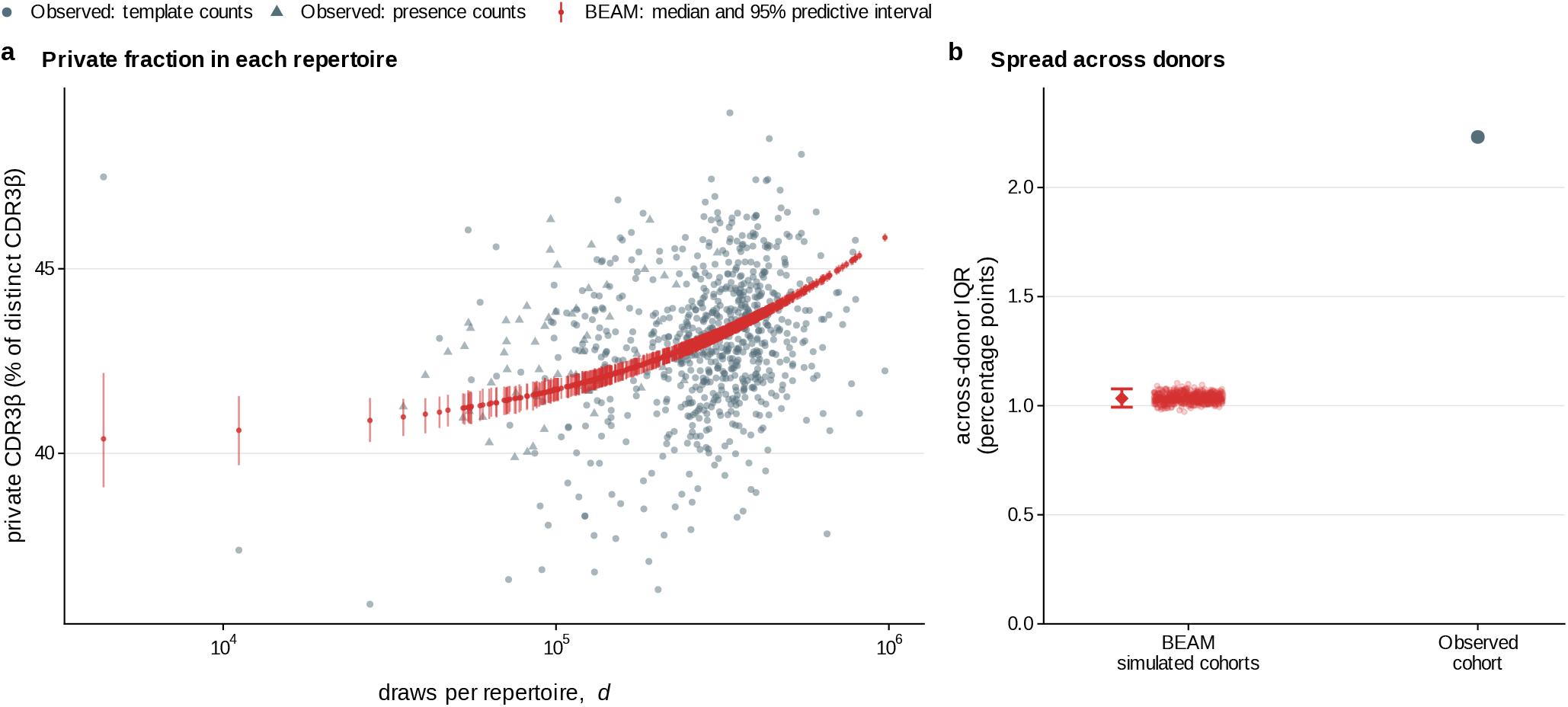
Predictive intervals for private fractions in held-out repertoires. All 768 Emerson donor repertoires [1] were excluded from model fitting. We sampled 512 independent complete cohorts from the fixed BEAM model, retaining each donor’s released count total: measured templates for 707 repertoires and one count per distinct amino-acid string for 61. For each donor, the private fraction is U/D, where U counts distinct CDR3*β* strings found only in that donor within the cohort and D counts all distinct CDR3*β* strings in that donor. Both counts vary across simulations. (**a**) Observed fractions are grey circles for measured-template repertoires and grey triangles for unit-presence repertoires. Red points and vertical segments show simulation medians and pointwise 95% predictive intervals at the original draw counts; 709 of 768 observed fractions lie outside their respective intervals. (**b**) Each small red point is the interquartile range (IQR) across donors in one simulated cohort. The adjacent red diamond and capped line show the median and 95% predictive interval of these IQRs; the grey point is the observed IQR. The observed IQR is 2.23 percentage points, compared with a simulated median of 1.03 and predictive interval of 0.99–1.08 percentage points. Sampling from the fixed shared model therefore does not reproduce the full observed variation across donors. Intervals describe sampling variation under the fixed model, excluding uncertainty in model fitting; donor intervals are not simultaneous. Private status is defined within the held-out cohort, whereas Fig. 4b uses the full release.

**Supplementary Figure S5:**
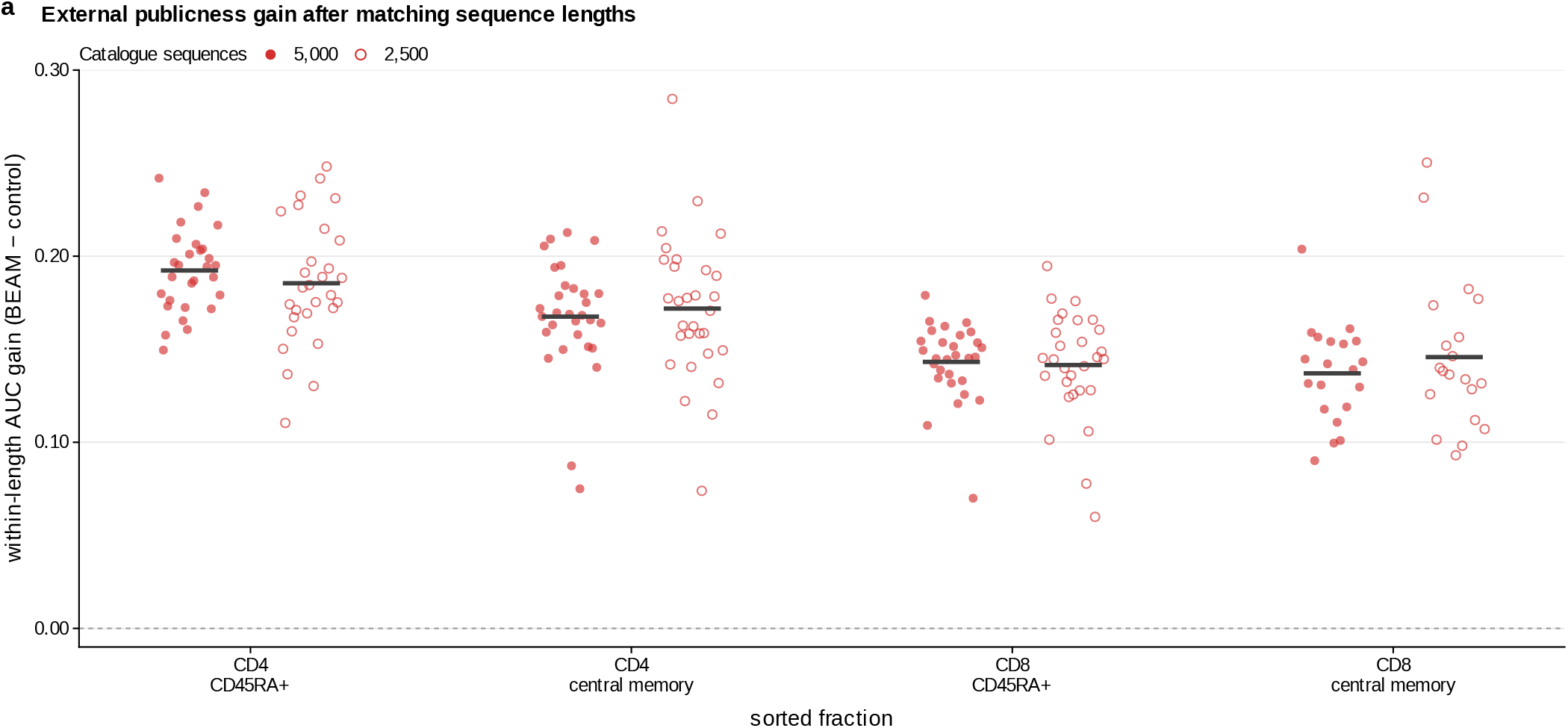
External publicness controls for length and catalogue size. This panel complements the primary ranking comparison in Figure 4h. 4 population-specific BEAM models trained on 10 Rubelt donors from 5 twin families [18] are evaluated in 30 donors from the separate ADC T1D dataset [20, 21]. Evaluated amino-acid sequences are absent from all training donors across all four populations. Public sequences occur in at least one other donor’s selected catalogue of the same sorted fraction; private sequences occur in none. The control Q is the same fitted model’s independent-position projection, preserving its length law and all individual-position amino-acid distributions conditional on length. (**a**) BEAM-minus-Q ROC AUC gain using only public/private sequence pairs of exactly equal length. Solid points use catalogues of 5,000 distinct amino-acid sequences sampled uniformly without replacement; open points use nested 2,500-sequence subsets. Shared/private labels are recomputed for each size. AUC pools tie-aware ranking wins over exact lengths, weighted by the number of public/private pairs. Each point is a donor/fraction comparison; horizontal offsets separate marks for display. Black bars average donors equally within each fraction and size; dashed line, zero gain. All 108 dependent donor/fraction gains are positive at both sizes. Fractions within a donor and sharing labels across donors are dependent; no inferential intervals or tests are shown. External CD8 CD45RA-positive fractions include naive and terminal effector-memory cells, and external central-memory fractions are narrower than the training memory labels. The endpoint is ranking within observed sequence catalogues; it does not test absolute sharing calibration or separate relationships associated with V/J usage. Importer-assigned record counts were not treated as molecular abundances.

**Supplementary Figure S6:**
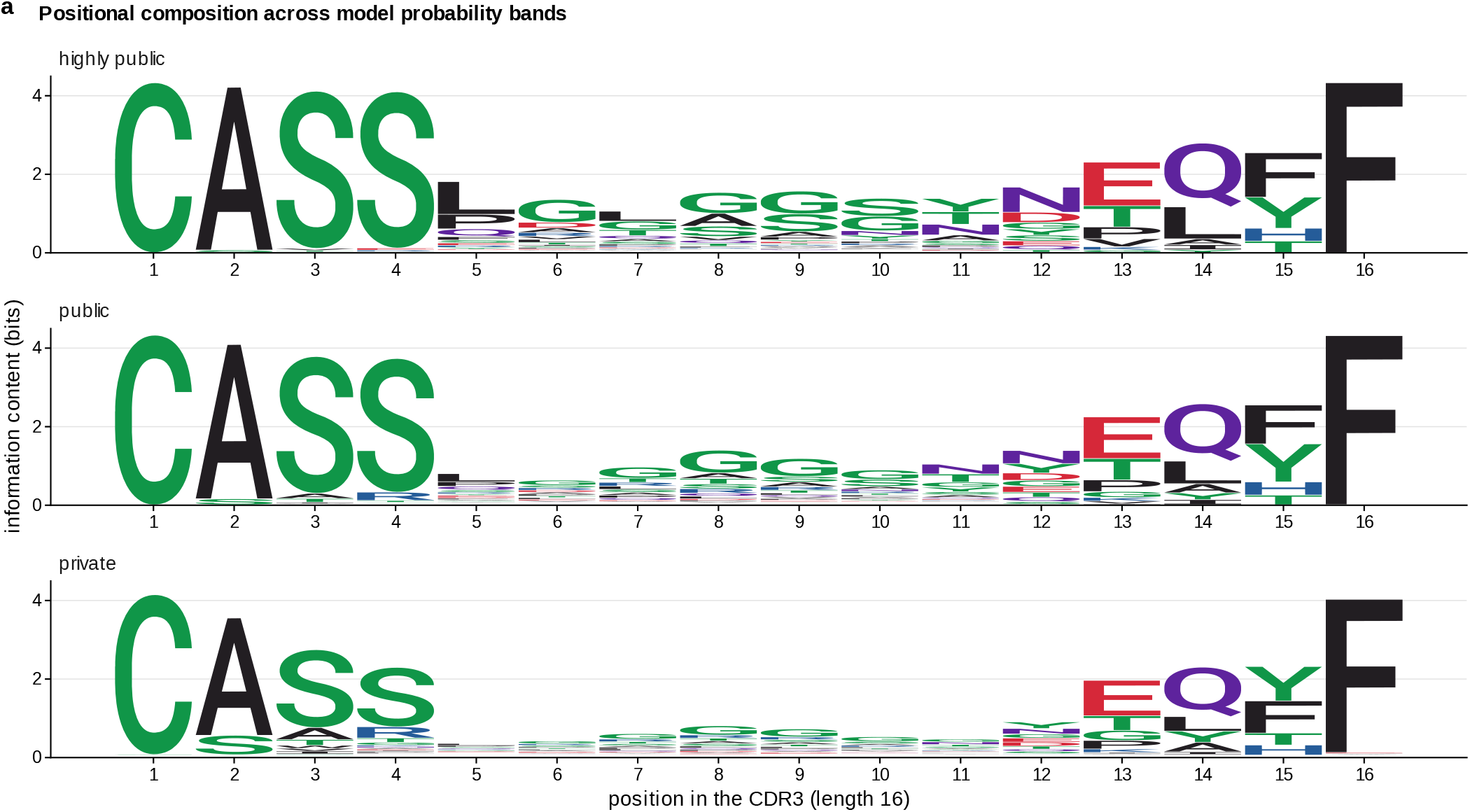
Sequence composition across model probability bands. (**a**) Probability-weighted sequence logos for length-16 CDR3*β* under the population reference model. Three disjoint bands of sequence observation probability *P*_seq_ are labelled by their expected publicness: highly public (above 10^−6^, corresponding to expected publicness about 4,520 at the boundary), public (between 3.55 *×* 10^−10^ and 10^−6^), and private (below 3.55 *×* 10^−10^). The lower boundary corresponds to expected occupancy in two repertoires at the original corpus depths. Within each band, sequences are weighted by their model probabilities, rather than counted equally. These are model-defined expected-occupancy bands, not observed public/private sequence groups. The three logos use a common information scale in bits; colours indicate amino-acid chemistry. Band-restricted values are calculated on a discretised probability grid. Positional consensus residues need not co-occur in a complete sequence within the corresponding band.

**Supplementary Figure S7:**
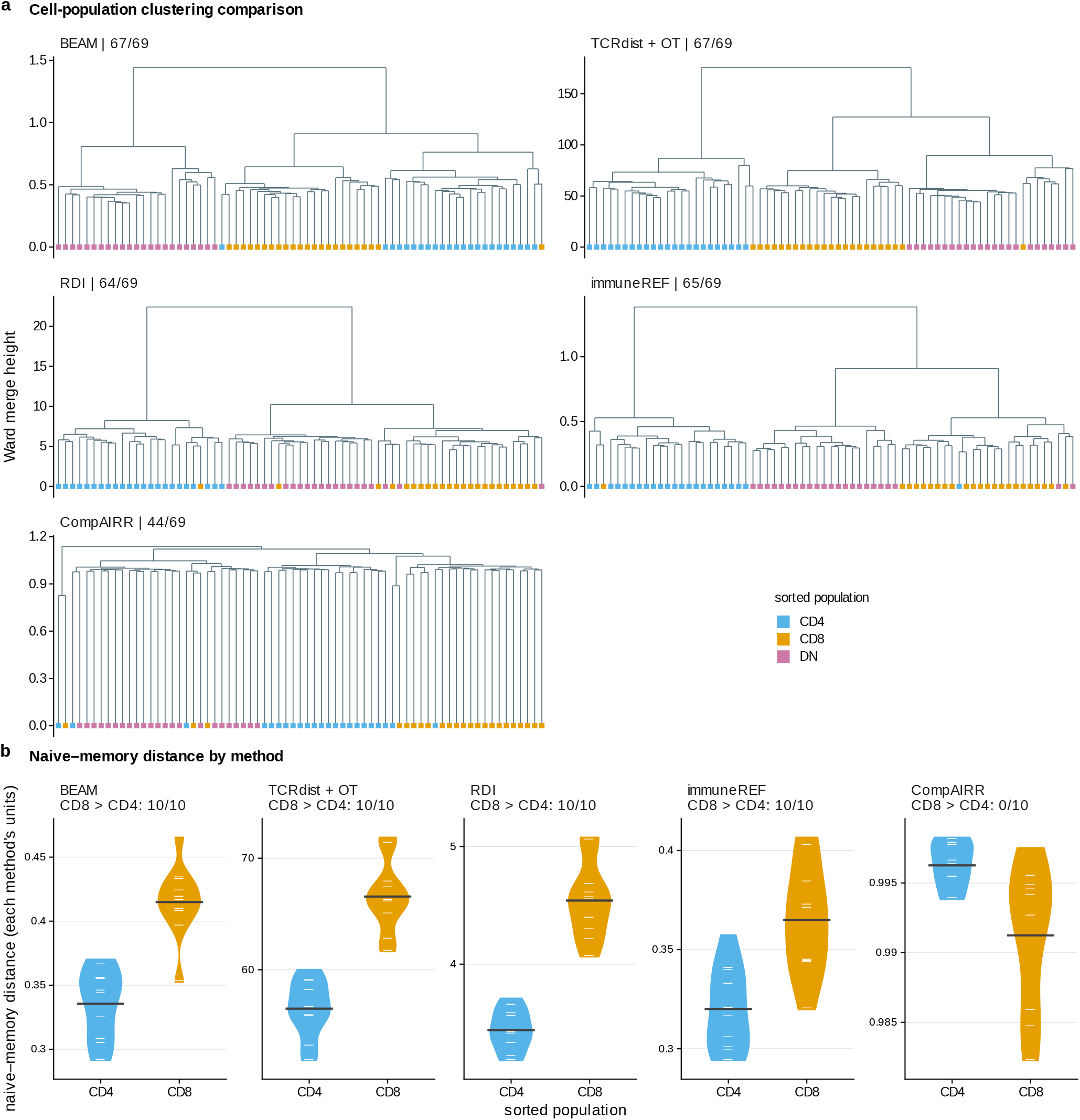
Repertoire distance methods compared on the benchmarks of Figure 5. (**a**) Ward dendrograms [27] of the same 69 mouse TRB repertoires [22, 17] (Fig. 5a) from 5 distance methods: BEAM distance; TCRdist with optimal transport [16, 17]; the Repertoire Dissimilarity Index [5] on V-gene calls; immuneREF [35] restricted to the four features available in these tables; and CompAIRR [39] exact amino-acid Jaccard distance (labelled BEAM, TCRdist + OT, RDI, immuneREF and CompAIRR). Each tree has its own merge-height scale; tips are coloured by sorted population (CD4, CD8, DN). The configurations and input representations of the methods are specified in Supplementary Methods. Strip labels give the number of repertoires placed with their population when each tree is cut into three clusters. (**b**) Distance between the naive and memory repertoires of each of 10 donors in 5 twin families (reannotated Rubelt cohort [18], as analysed by Bolen et al. [5]; Fig. 5b), in CD4 and CD8, from the same five methods, one panel per method on its own scale. Violins, donors; white ticks, individual donors; horizontal bars, means. Strip labels give the number of donors whose CD8 distance exceeds their CD4 distance. Concordance and donor ordering are descriptive comparisons on these datasets.

**Supplementary Figure S8:**
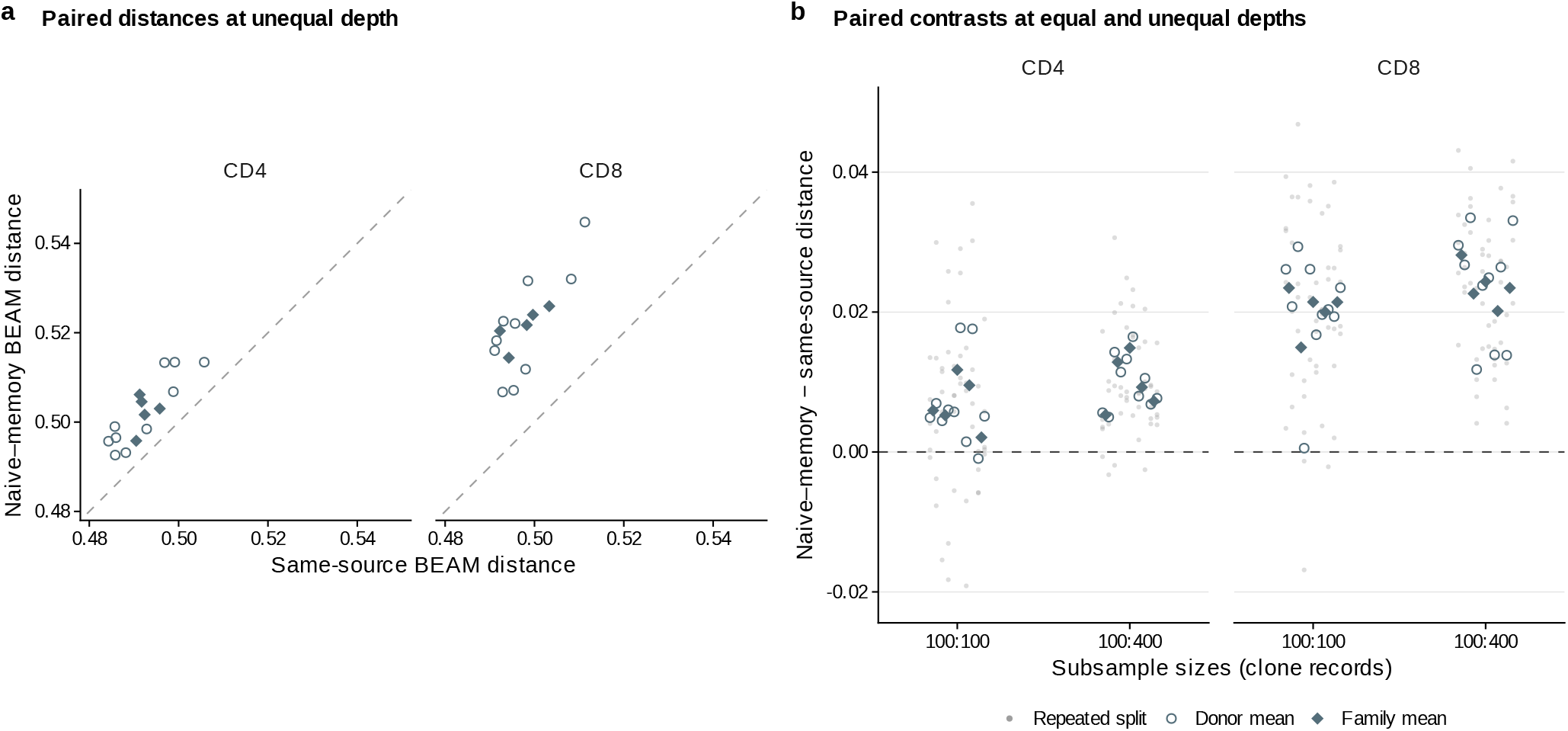
Paired repertoire contrasts under unequal sampling. Reannotated Rubelt twin-cohort repertoires [18] are the same inputs as Fig. 5b/c. The same 10 donors in 5 twin families contribute CD4 and CD8 repertoires at 100:100 and 100:400 clone records, with 5 assignments per donor and condition. Disjoint subsets of each observed source table are fitted independently. Same-source distance averages naive–naive and memory–memory comparisons; naive–memory distance averages both assignments of phenotype to the smaller sample. Each comparison uses the stated pair of depths. (**a**) Raw distances at 100:400 records. Open circles, donor means; diamonds, twin-family means. The diagonal denotes equal naive– memory and same-source distances; both facets use the same square scale. (**b**) Naive–memory minus same-source distance at both depth conditions. Faint dots retain individual assignments, including negative contrasts; circles and diamonds are as in a. Dashed line, zero. At unequal depth, all 10 CD4 and 10 CD8 donor-mean contrasts are positive; 46 of 50 CD4 and 50 of 50 CD8 assignments have positive averaged contrasts. This average does not imply a positive contrast in each orientation: CD4 donor means are positive in 10 of 10 donors when naive is smaller and 7 of 10 when memory is smaller. Repeated assignments and twins are dependent; no inferential intervals or tests are shown. This follow-up uses previously specified splits from the existing cohort, not independent assay replicates or prospective validation. The plotted contrast is not an unbiased population-distance correction; raw distances remain depth dependent, and no superiority over other methods is established.

**Supplementary Figure S9:**
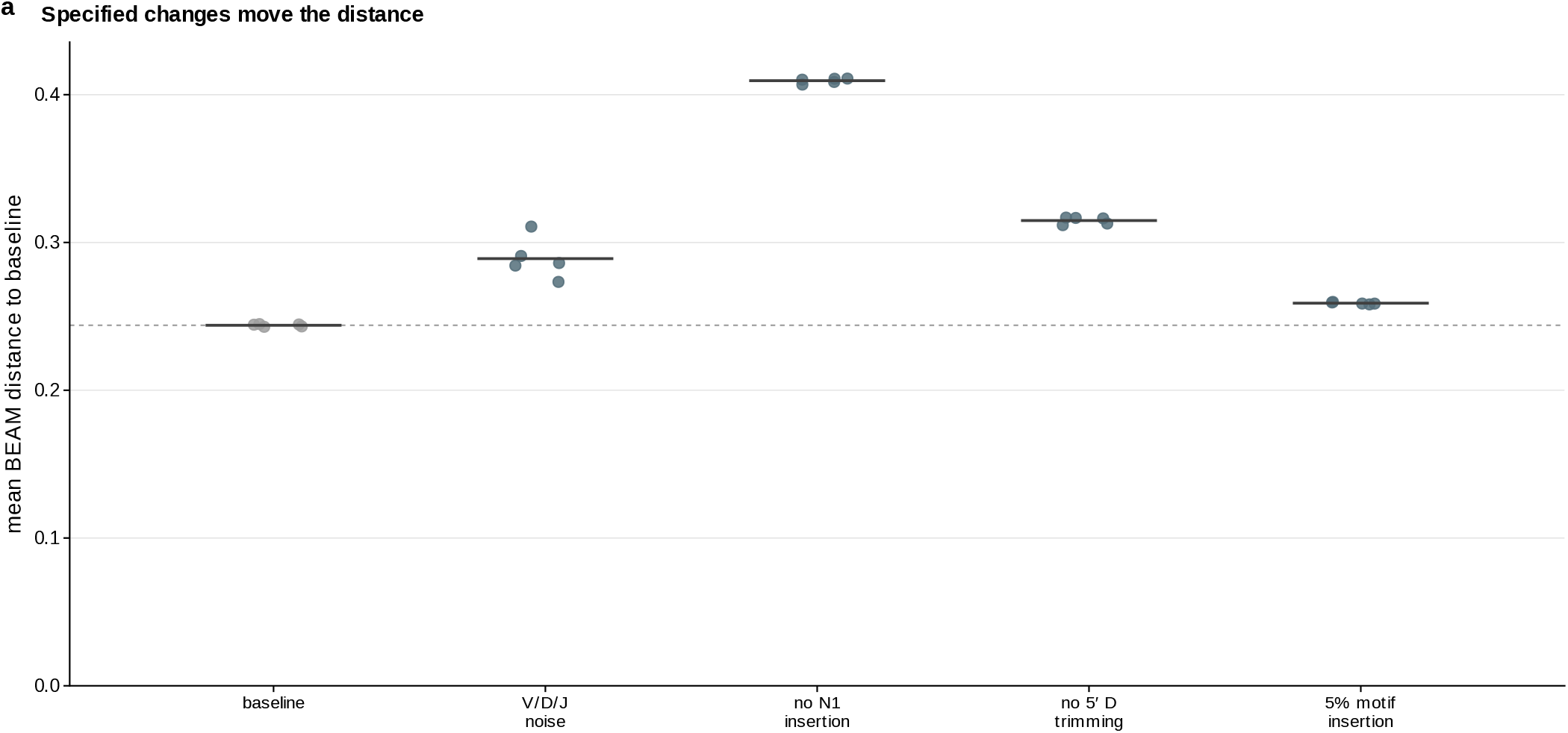
BEAM distance under controlled repertoire perturbations. (**a**) Mean BEAM distance to baseline repertoires in an adaptation of the controlled-perturbation benchmark of immuneREF [35], with repertoires simulated by immuneSIM [23]. The 25 human TRB simulations comprise 5 independently generated repertoires per condition, each containing 10,000 retained amino-acid sequence records with unit weight per record. Conditions are default generation, V/D/J usage noise, removal of N1 insertion at the V–D junction, removal of 5^*′*^ D trimming and nominal 5% motif implantation (YAY or YVY). Points, repertoires; horizontal bars, condition means. Each point averages distances to all baseline repertoires, excluding itself when applicable. The dashed line is the mean distance among distinct baseline replicates. One-sided exact repertoire-label permutation tests compare mean between-condition distance with the average of the two within-condition means; they do not test only the plotted height above the dashed line. Holm adjustment [38] covers all 4 perturbations: adjusted *P* = 0.0317 for no N1 insertion, 0.0317 for no 5^*′*^ D trimming, 0.0317 for motif implantation and 0.357 for V/D/J noise. These simulations assess sensitivity to specified sequence changes.

**Supplementary Figure S10:**
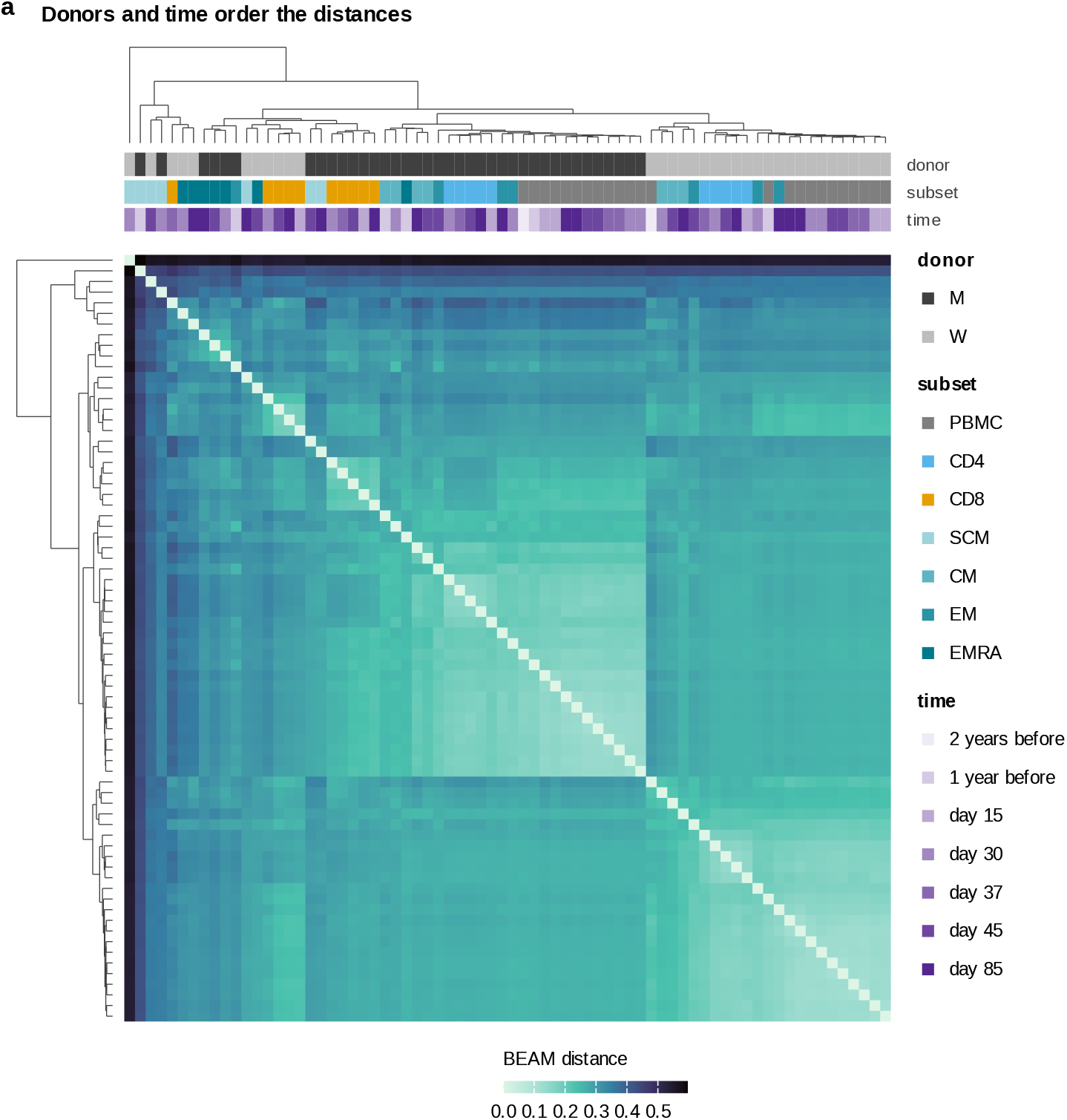
Repertoire distances between two people over time. (**a**) BEAM distances between 72 TCR*β* repertoires represented by selected top-clone records from two donors sampled over time (Minervina et al. [24]; 38 repertoires of donor M and 34 of donor W), ordered by complete-linkage clustering of Euclidean distances between distancematrix row profiles (trees, top and left). Strips: donor, cell subset and time point. PBMC, peripheral blood mononuclear cells; SCM, stem cell-like memory; CM, central memory; EM, effector memory; EMRA, effector memory with CD45RA re-expression. Day 0 is return from travel; symptoms began on day 3. Earlier samples were collected one or two years before infection. One BEAM model is built for each repertoire, with equal weight per clone record; their local-block Hellinger distances compare length and local sequence distributions (Methods). The two-donor comparison is descriptive.

**Supplementary Figure S11:**
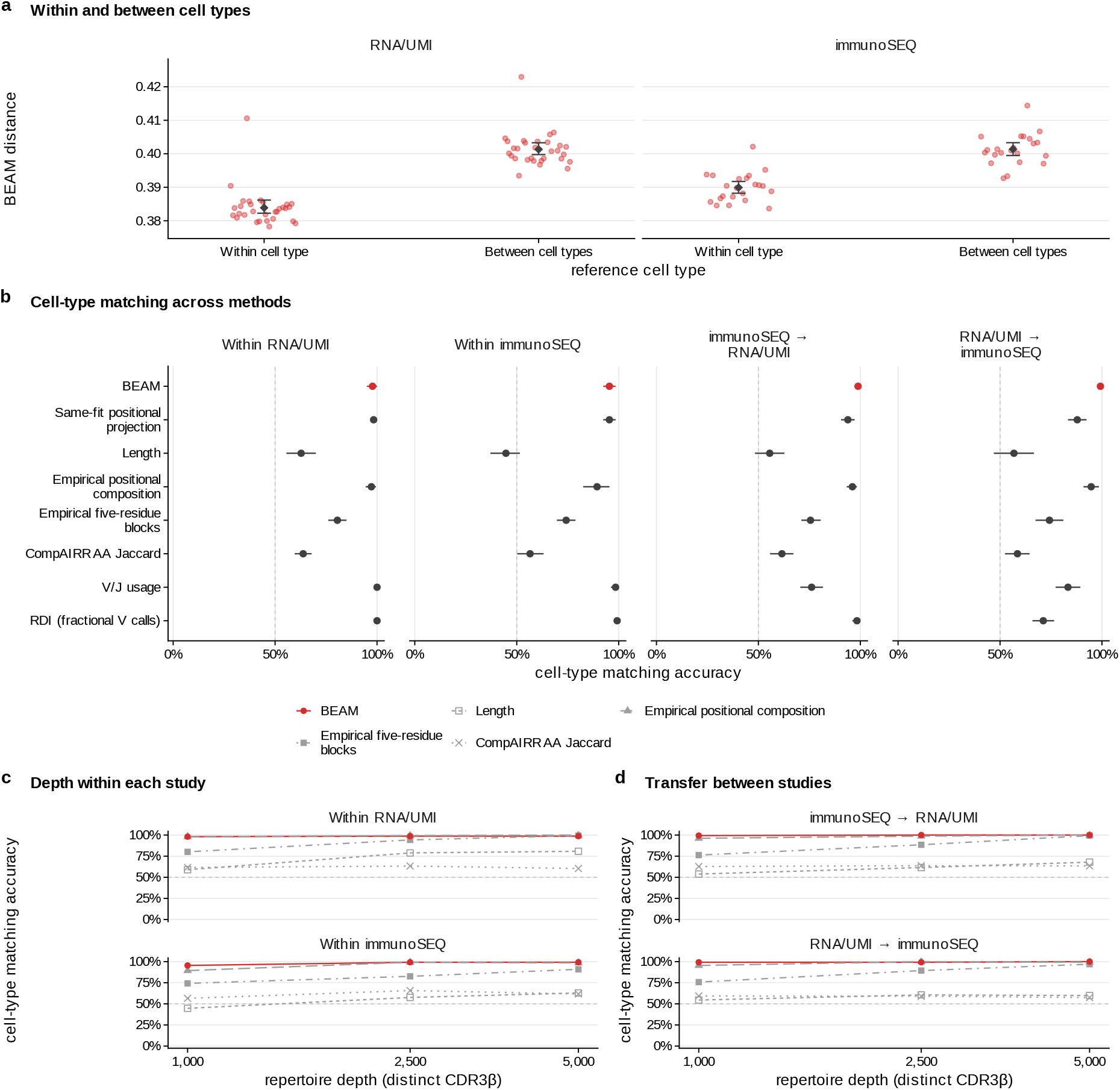
Human CD4 and CD8 repertoire matching across studies and input depths. Paired peripheral-blood CD4/CD8 repertoires comprise 30 GSE285951 donors (McCord et al. [25]; RNA with unique molecular identifiers, RNA/UMI) and 22 immunoSEQ donors (Emerson et al. [26]). GSE285951 is external; Emerson repertoires were represented in the corpus. Models are fitted separately to each repertoire, without a population reference model, using the stated number of distinct standard-amino-acid CDR3*β* strings (length ≤27), each weighted once. (**a**) At depth 1,000, red points show donor means of BEAM distances to other donors’ same-type or opposite-type repertoires, averaging CD4/CD8 queries and 3 subsamples. Black diamonds and whiskers, donor means and 95% confidence intervals. Points are offset; facets share a nonzero-origin axis. Within/between means are 0.384/0.401 (RNA/UMI) and 0.390/0.401 (immunoSEQ). (**b**) Nearest-neighbour matching accuracy and 95% confidence intervals for all predefined methods at depth 1,000. Within-study references exclude both query-donor repertoires; sequences need not be unseen. Arrows, reference *→* query; no target-label calibration. BEAM is red; controls, grey. The same-fit positional projection retains BEAM’s length and single-position marginals (also Fig. 5d); empirical positional composition is a separate control. (**c, d**) Within-study matching (c) and reciprocal transfer (d) across depths in the same 26 RNA/UMI and 22 immunoSEQ donors. Shapes, methods; lines join measured depths; whiskers, BEAM confidence intervals. Complete tables retain all intervals and the higher-depth extension. Dashed lines in b–d, 50% chance for balanced classes. Subsamples are nested within 3 repetitions. Repetition and CD4/CD8 scores are averaged within donors before 10,000 donor-bootstrap draws; intervals condition on the observed reference cohort. V/J and RDI [5] also use gene annotations, with fractional weighting of ambiguous/unassigned calls. Transfer combines study, assay and clinical-cohort differences; it does not isolate assay effects or test disease/naive–memory classification. Cohort composition, annotation completeness and full procedures are given in Supplementary Methods.

**Supplementary Figure S12:**
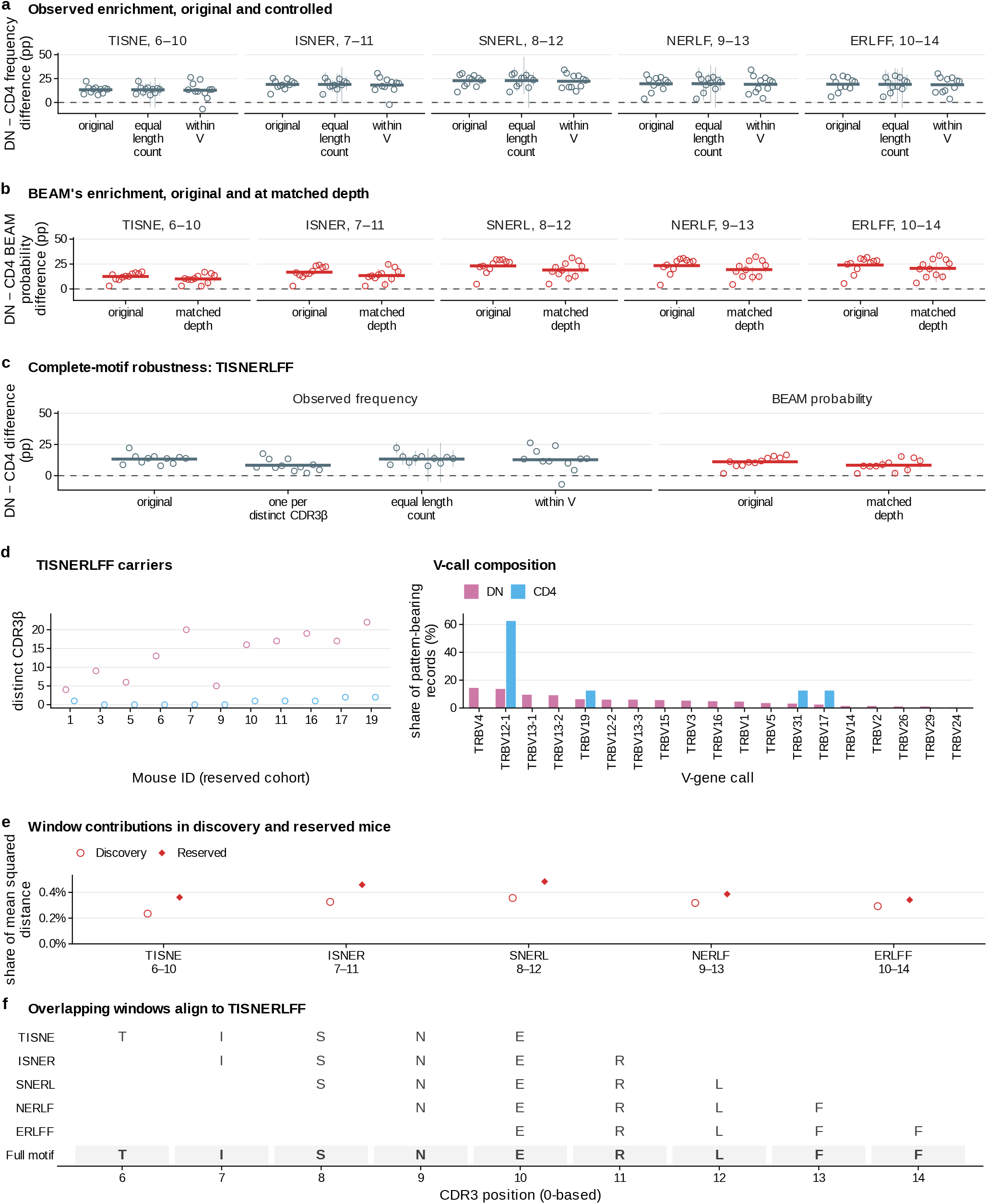
Window contributions, controls, alignment and receptor context. Panels a–d use the 11 reserved mice [22, 17] of Fig. 5f. (**a, b**) Columns show the five selected windows, with zero-based positions; their literal overlap is aligned in f. Values are DN − CD4 differences conditional on length 15, in percentage points. **a**, observed clone-record frequencies: original; equal length count, sampling both populations without replacement to the smaller length-15 count; within V, pooled-count-weighted differences across V-call strata shared by both populations. Equal-count points average 200 draws; segments span their 2.5th–97.5th percentiles. **b**, exact BEAM probabilities from original models or after matching total record counts. Matched-depth points average 3 paired refits; segments span their minimum and maximum. (**c**) Complete TISNERLFF motif at positions 6–14, including a control counting each distinct CDR3*β* once per repertoire. Observed matches coincide with TISNE; model probabilities are queried separately. Distinct-CDR3 and within-V differences are positive in 11 and 10 mice; matched-depth mean model contrasts are positive in 11. In a–c, points are mice, horizontal bars are unweighted mouse means and dashed lines mark zero. Intervals describe subsampling/refit variation, not confidence intervals across mice. (**d**) Left, distinct motif-bearing CDR3*β* per mouse and population, including zeros; x-axis numbers identify the reserved mice. Right, V-call frequencies among motif-bearing records pooled across mice, normalised separately for DN (284 records) and CD4 (8). Pooled DN contains 43 distinct CDR3*β* and 19 V calls; the most frequent V call contributes 14.4% of records. Distinct counts deduplicate across mice. V stratification is observational. (**e**) Window contributions in 12 discovery mice (open circles) and 11 reserved mice (filled diamonds). Each share is mean window contribution divided by mean paired D^2^, not the mean per-mouse share. Discovery contributions alone selected these windows. Summed shares are 1.53% and 2.03%, respectively. These sum original window terms, not a new complete-motif term; no uncertainty intervals are shown. (**f** ) The five windows aligned by their zero-based coordinates, with their complete TISNERLFF span at positions 6–14 shaded below. The complete-event controls and carriers are evaluated separately in c and d. Grouping compatible overlaps was post hoc. These windows are related terms in one group, not independent motif families. The follow-up uses previously examined mice and does not establish antigen specificity.

## Notes

### Competing Interest Statement

GY is a scientific advisor of Clareo Biosciences.

